# Assembling Neural Latching Switch Circuits for temporally structured behavior

**DOI:** 10.64898/2026.04.15.718666

**Authors:** Alexis Dubreuil

## Abstract

Elaborated temporal structures in behavior have been recognized as a hall-mark of high-level cognitive processes, such as planning or natural language. Systems neuroscience experiments in behaving animals have validated cell assemblies as a fundamental piece of neural circuitry underlying lower-level cognitive processes. Although recent research has identified neural bases for certain temporally structured behaviors, it is still an open question whether a general-purpose circuit - akin to cell assemblies - supports such behaviors. Here we show how Neural Latching Switch Circuits can be assembled as lego-bricks to create neural architectures suited for the implementation of *any* procedural behavior. By focusing on various forms of temporal structures and leveraging a combination of theoretical tools, we demonstrate how these circuits can be mapped onto interacting brain areas and propose predictions for identifying analogous structures in mammalian brains, for instance in cortical columns. By interpreting neural architectures as computing automata we reveal surprising relationships between behavior and computation.

## Introduction

Early works in psychology and neuroscience recognized that behavior, and especially high-level cognitive processes such as planning, language or reasoning rely on temporally structured sequences of mental events (1). These works were accompanied by speculations about potential underlying neural substrates, but these remained vague given the amount of experimental knowledge available at the time (2). Looking to propose such a neural substrate, one is facing two challenges.

The first one is to define relevant temporal structures to model behavior. Temporal structures have been introduced to model animal behavior under laboratory conditions, such as the swimming patterns of zebrafish larvae (3), locomotion of C. Elegans (4), courtship songs of birds (5) or grooming patterns of rodents (6). Another more systematic attempt of such a definition is Chomsky’s generative grammar (7) which defined classes of languages organized in a hierarchy, with qualitatively different forms of temporal structure at each complexity level (8). However, the relevance of this classification for modeling behavior has been questioned (9; 10; 11).

A second challenge is to relate temporal structures to the anatomical and physiological properties of brain circuits. Oscillations in electrical activity for instance have been deemed important to support cognition (12). Another set of experimental results has demonstrated that the concept of attractor neural network is well suited to account for the emergence of elementary cognitive processes involved in working memory, spatial navigation or decision making (13). They formalize Hebb’s ideas (14), with a synaptic connectivity structure characterized by positive feedback loops, allowing electrical activity to build-up among neurons. This leads to the activation of groups of neurons, or cell assemblies, that encode behavioral variables such as the identity of a remembered image (15), a location in space (16) or a choice option (17). These attracting neural states in principle persist over arbitrarily long durations and therefore do not seem well suited to account for the emergence of high-level, temporally structured, cognitive processes. Systems neuroscience studies have focused on specific temporally structured behaviors (see e.g. (4; 6; 18; 19; 20; 21)), but whether or not a generic circuit motif, akin to cell assemblies, sustains such behaviors remains to be explored (13; 22).

In order to address these two challenges, we focus on a set of tasks exhibiting primitive temporal structures which have been proposed to underlie behavior (23). We show how Neural Latching Switch Circuits (**NLSC**), attractor neural networks augmented with gate neurons (24), can be assembled as lego-bricks into neural architectures solving all of these tasks. Using models of binary neurons, we compare the efficiency of these architectures with alternative ones. We outline the coding and structural properties of these architectures, and discuss how those are related to physiological and anatomical characterizations derived from systems neuroscience experiments. This allows us to address the second challenge and to propose that interacting NLSC can be used to model interactions between cortical columns. To describe the relationship between neural architectures and the temporal structures they support, cell assembly activations are naturally interpreted as automata states (25), with different forms of temporal structures mapping onto different automata abilities. Picturing neural networks as computing machines manipulating symbols leads us to outline temporal structures fundamental to behavior. This addresses the first challenge and gives a computational justification for the taxonomy of temporal structures proposed in (23).

### 1 External memory for behaviors with long-distance dependencies

In previous work we introduced Neural Latching Switch Circuits (24). They are composed of an attractor neural network, implementing a dictionary of stable states, together with gate neurons, allowing to program transitions between states upon presentation of external inputs. We showed that NLSC can implement simple behaviors involving sensory processing or motor generation. For instance NLSC can be mapped onto the fly’s head direction system (26). Or, more generically, they can be mapped onto finite-state automata (**FSA**, Method 6.5). FSA are abstract models of computation with restricted computational abilities (25). Within the framework of generative grammar, FSA can only recognize languages that exhibit a simple form of temporal structure, corresponding to the lowest class of Chomsky’s hierarchy (SI Fig. 1).

To depart from this simple form of temporal structure, we considered long-distance dependencies which are commonly described in behavior. An example is subject-verb agreement in sentences where the subject is separated from the verb by other words. Another example is canari’s courtship songs where the probability to produce a phrase depends not only on the currently sung phrase, but also on preceding phrases sung several steps in the past (5). Following what we previously did to model canari’s song production (27), we trained artificial neural networks (**ANN**) on a minimal task involving long-distance dependencies (Fig. 1a, Method 5.3.2). Networks are presented with one of two initializing cues and are asked to produce one of two sequences that share a common path of production states, and then diverge. Training a NLSC with a 5-states dictionary (Method 5.2.1), one per symbol to be produced, did not allow to implement the task, but augmenting the architecture with extra neurons we obtained good performance.

**Fig. 1:**
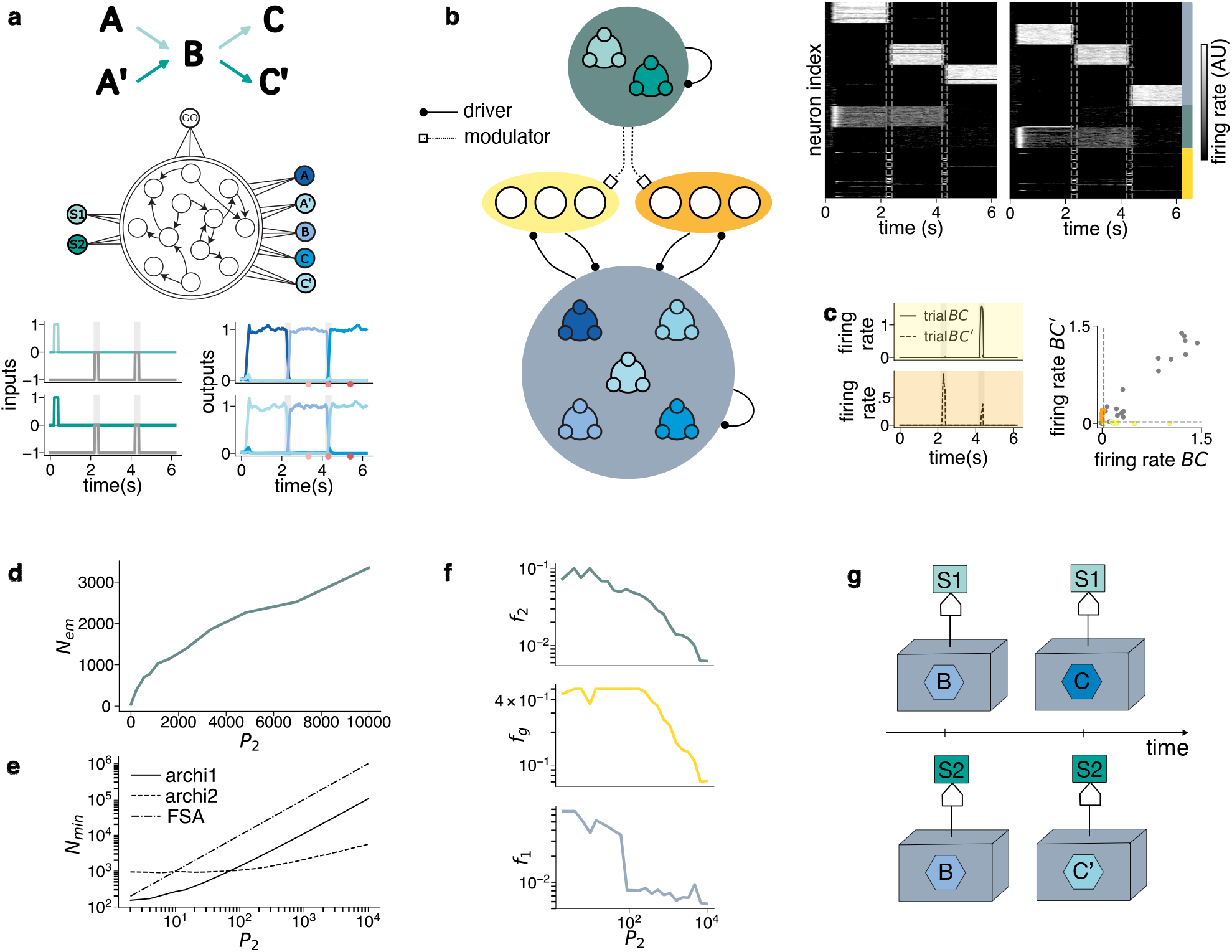
NLSC augmented with an external memory. **a**, ANN trained to produce a minimal set of two sequences S1 and S2 with long-distance dependencies. Bottom: inputs and trained outputs in the two different types of trials. Grey shades represent presentation of the go input releasing inhibition onto gate neurons. **b**, Left: Schematic of a NLSC, blue dictionary and yellow/orange gate neurons, augmented with an external memory (green). The dictionary is pictured with 5 cell assemblies, the external memory with 2 cell assemblies. Each of the two external memory states controls a permutation of dictionary neurons, by controlling the gain of one of the population of gate neurons. Right: Raster of neural activity underlying the production of the two sequences. **c**, Left: Firing rate of two exemple gate neurons driving transitions from B to C (top) and from B to C’ (bottom). Right: Firing rate of all gate neurons in transition B to C (X-axis) and B to C’ (Y-axis). The most selective gate neurons, controlling transitions, are colored (Method 5.3.2). **d**, Number of neurons in the external memory for optimized architecture. **e**, N_min_ for various neural architectures implementing a set of P_2_ sequences on a dictionary of P_1_ = 30 states. **f**, Coding levels in external memory (top), gates (middle) and dictionary (bottom) for the optimized architecture. **g**, Time evolution of a FSA (light blue box) augmented with an external memory (green square) during the production of S1 and S2. The front-side of the box displays the state in which the automaton is in.

In such an architecture the NLSC is augmented with an external memory that maintains a contextual neural representation of the initializing cue (Fig. 1b), allowing to bias the transitions towards the correct dictionary state at the point where the two sequences diverge. This routing is accomplished by a specific connectivity from the external memory to the gate neurons of the NLSC. Gate neurons exhibit non-linear mixed-selectivity (28), activating upon conjunction of the go cue and one of the contextual neural representations (Fig. 1c, SI Fig. 3a,d). In SI Fig. 4, we describe the functioning of this neural architecture at the population level, showing how the state of the external memory modulates the gain of two populations of gate neurons, controlling interactions between cell assemblies to correctly route neural activity (Method 5.2.3).

In order to provide a more quantitative description, we embodied this type of neural architectures in models of binary neurons, and adapted storage capacity calculations developed for studying attractor neural networks or perceptrons with binary synaptic weights (29; 30; 31). In the spirit of (32; 33) these models are optimized by finding the coding and structural parameters that minimize the required number of neurons, *N*_*min*_ (see Method 5.5 or (24)). We computed *N*_*min*_ and analyzed the characteristics of neural architectures composed of a NLSC with *P*_1_ dictionary states and an external memory with *P*_2_ states (SI Section 6.1.3). Sparse but distributed representations of the *P*_2_ external memory states are stored as stable states in an attractor neural network whose size *N*_*em*_ scales as 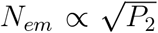 (Fig. 1d). As for the case of FSA (24), using a NLSC with *P*_2_ gate populations was more efficient for *P*_2_ *P*_1_ (Fig. 1e, full line), compared to using a NLSC with *P*_1_ gate populations (Fig. 1e, dashed line). The converse being true for *P*_2_ ≫ *P*_1_. Such an architecture storing *P*_2_ = 10, 000 contextual representations, here thought of as words piloting a dictionary of phonemes of size *P*_1_ = 30, requires *N*_*min*_ ≃ 6, 000 neurons. Neurons inside a cortical column could thus implement an articulatory apparatus mapping word representations into their motor production, with sparse neural patterns in external memory, dictionary and gate populations (Fig. 1f).

From a computational point of view, this neural architecture is interpreted as a finite-state automaton augmented with an external memory (Fig. 1g): the NLSC implements a FSA (24), with state transitions in response to the go cue that are determined by the state of the external memory. The task above (Fig. 1a) involves a finite number of sequences to be produced and could in principle be implemented by a FSA and thus a NLSC (24), without resorting to an external memory. By training ANN with different constraints (Method 5.2.2) we found other network mechanisms that could support the production of sequences with long-distance dependencies (SI Section 5.3.2). A different mechanism is one where symbol B is produced by different neural states depending on sequence identity (SI Fig. 3k). From a computational point of view, such an implementation is interpreted as a FSA, without any external memory, with distinct states associated to the production of the same symbol in different contexts (SI Fig. 3l). Computing *N*_*min*_ for this neural architecture allows to show it is not advantageous compared to the architecture with external memory we presented first (Fig. 1e, dotted-dashed line, SI Section 6.1.3).

Long-distance dependencies are present in various behavioral tasks used in neuroscience experiments, neural representations for dictionary (34; 35) or external memory (36; 37) states have been exhibited in different cortical networks. It would be interesting to see whether gate neurons mediating communication between cortical areas could be described.

### 2 NLSC as a controllable external memory for behaviors organized into chunks

Behavior can often be described as hierarchically organized. For example, the different locomotive behaviors of zebrafish larvae, such as during hunting or counter-stream swimming, can be decomposed into temporal sequences of thirteen elementary tail movements, which are themselves a sequential combination of muscle activations (3). Here we assemble NLSC (Method 5.2.4) to build a neural architecture that implements such a motor hierarchy, producing sequences organized into chunks (23) (Fig. 2a, Method 5.3.3). To do so we built upon the architecture above (Fig. 1b) by adding an extra state in the dictionary, to be played at the end of each chunk (Fig. 2b, light blue cell assembly). This stop state triggered transitions in the external memory through a new set of gate neurons (Fig. 2b, orange), which are bidirectionally coupled with the external memory and that receive connections from the low-level dictionary (Fig. 2b, blue). This leads them to exhibit the following non-linear mixed selectivity: neurons respond to conjunction of the middle-level dictionary states and the added stop state (SI Fig. 5). Thus coupling two NLSC allowed to produce sequences organized into chunks (Fig. 2c).

**Fig. 2:**
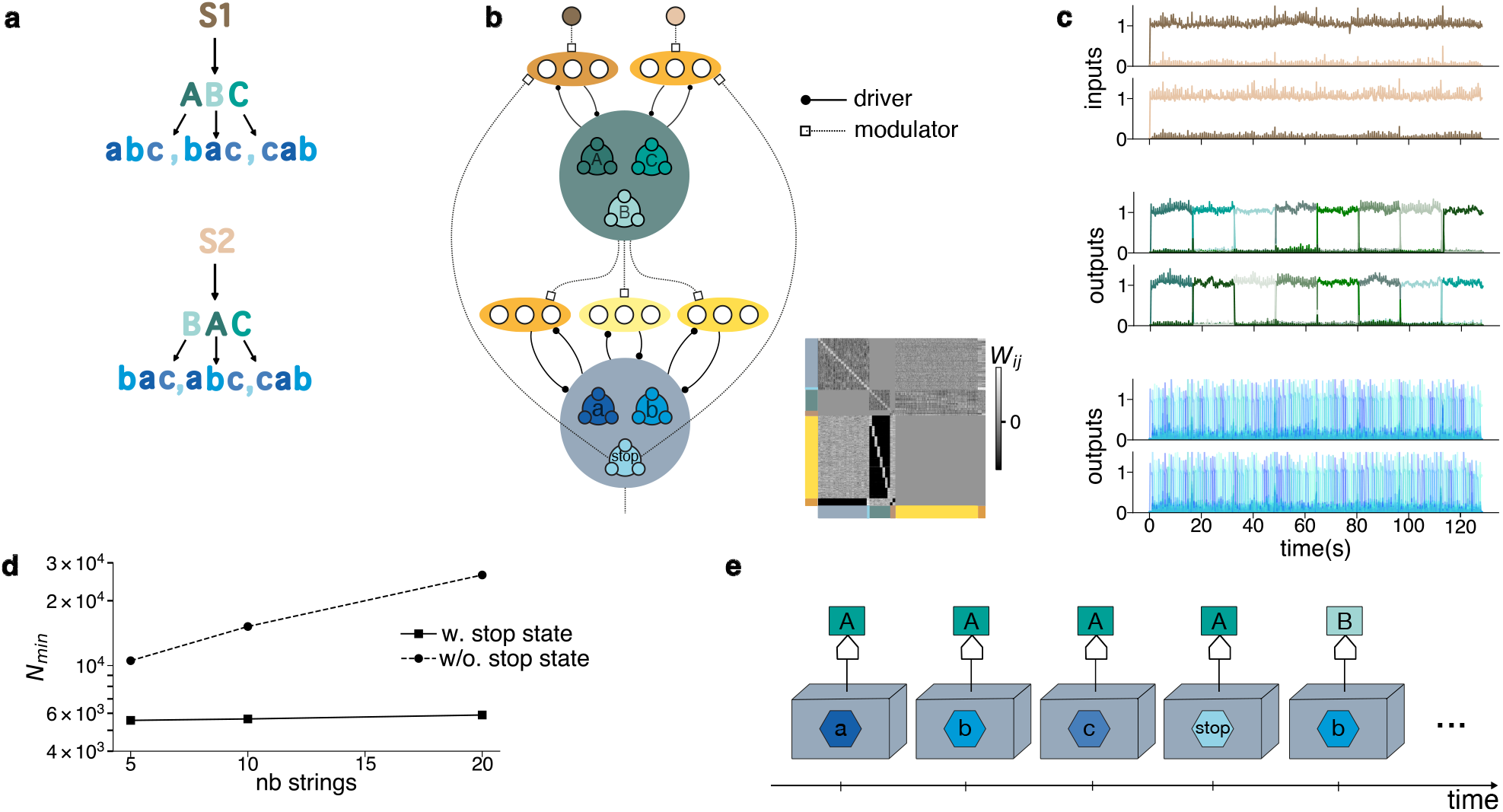
Assembling NLSC for behaviors organized into chunks. **a**, Temporal structure of sequences organized into chunks. **b**, Coupling two NLSC. Green cell assemblies represent chunk identity, blue cell assemblies represent terminal symbols to be produced. Gate neurons (orange shades) program transitions between cell assembly activations. **c**, Cell assembly activations in the different levels of the architecture during the production of S1 (top) and S2 (bottom). Each of these sentence is composed of 8 chunks (middle, green). At the lowest level, each chunk is composed of a sequence associated to a permutation of 20 states (bottom, blue) **d**, N_min_ for neural architectures with and without stop states in the low-level NLSC. **e**, Time evolution of an automaton summarizing the behavior of the proposed neural architecture. A FSA reads from the external memory to determine its next state. When in the stop state, the FSA writes a new symbol on the external memory, programming a new set of transitions in the FSA.

By computing *N*_*min*_ for architectures with and without stop states (SI Section 6.1.4), we found that using a stop state is beneficial: for a motor apparatus wired to produce 10, 000 words and 5 sentences of 100 words, the number of required neurons goes from *N*_*min*_ = 5, 500 with a stop state to *N*_*min*_ = 10, 500 without. This advantage becomes even more pronounced when the number of sentences to be produced increases. For 20 sentences, it goes from *N*_*min*_ = 5, 900 with a stop state to *N*_*min*_ = 26, 300 without (Fig. 2d).

Such interacting NLSC share similarities with the large-scale organization of motor cortex. Human motor cortex has been described as an intertwining of functionally low-level motor-effectors regions with functionally higher-level regions (38). Accordingly, electrical stimulations in monkeys’ motor cortex showed that at some location, brief stimulations elicited activation of single motor effectors such as finger moves. By contrast, in nearby regions, brief stimulations were inoperant but more prolonged ones triggered the execution of whole motor programs such as moving upper limbs to direct the hand towards the mouth (39). We expect neural activity in the high-level regions to evolve more slowly than in the low-level ones (40) (Fig. 2c). This can be tested using recording methods with high temporal resolutions, allowing to characterize the time-scales of the evolution of neural activity in the different regions of motor cortex during complex movements.

The automaton corresponding to the two coupled NLSC (Fig. 2e) has an extra feature compared to the one we proposed for the minimal task with long-distance dependencies (Fig. 1g): machine states can now update the state of, or write on, the external memory. Writing on the external memory corresponds to the low-level dictionary signaling the upper-level dictionary of its current state, and triggering transitions when the low-level state goes out of the null-space of the high-level NLSC (41).

### 3 NLSC as an attentional focus for behaviors with algebraic patterns

Another form of temporal structure that has been highlighted by cognitive neuroscience is algebraic patterns (23), characterized by the ability to recognize or produce patterned data, e.g. XYX, where X and Y are roles that can be filled with symbols from an alphabet (Fig. 3a, Method 5.3.4). In order to assemble an architecture suitable for such a production, we took inspiration from recordings of neural activity in prefrontal cortex while monkeys memorize and produce temporal sequences XYZ of three roles, with an alphabet of fillers constituted of eight spatial locations (18). The neural patterns associated to the memory of spatial locations for each role were carried by distinct groups of neurons (18; 42). In Fig. 3b,c we present a neural architecture producing temporal sequences with an algebraic pattern structure. Two dictionaries (green) form an external memory and are used to store a representation of the content of roles X and Y, they project to neurons responsible for the production of the symbols (blue). Interactions between the external memory and the production network are dynamically modulated by a NLSC responsible for producing the abstract sequence XYX (purple). Each dictionary state of this NLSC couples one of the external memory cell with the production network, or machine circuit. Modulation of couplings are performed through a new type of *output* gate neurons associated with dictionaries of the external memory (orange). As opposed to the *transition* gate neurons that we have discussed thus far, they do not feedback to their dictionaries.

**Fig. 3:**
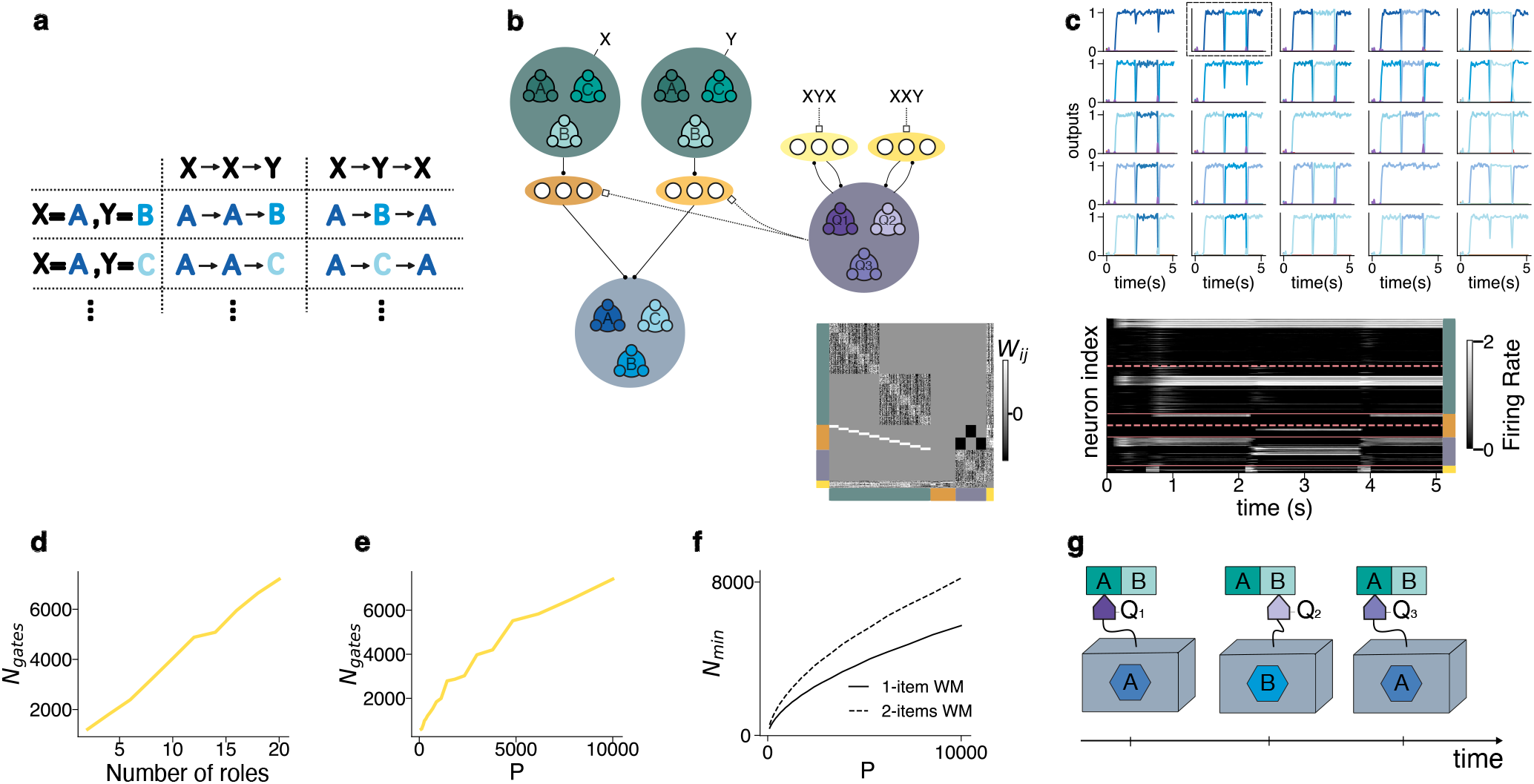
Assembling NLSC for behaviors with algebraic patterns. **a**, Sequences for two algebraic patterns XXY and XYY. Each role X or Y can be filled with a symbol A, …, Z. **b**, Neural architecture for producing sequences with an algebraic patterns temporal structure. An output dictionary (blue), reads the state of one of two dictionaries playing the role of two external memory cells. The current dictionary state of the head NLSC (purple) decides from which cell the output is read, trough modulation of populations of output gate neurons (orange). **c**, Top: readout activations (blue cell assemblies) for a network assembled to produce sequences XYX with 5 possible different fillers. Bottom: raster of neural activations in the purple and green NLSC during the production of sequence ABA, cf dashed square on Top. **d**, Number of output gate neurons for an optimized architecture as a function of the number of slots in the external memory, and **e**, the number of possible symbol in each memory slot. **f**, Minimal number of dictionary neurons for an external memory implemented by a single attractor network operating in a two-items working memory regime (dashed) or two attractor networks with single item working memory (full). **g**, Automaton interpretation of the neural architecture. A head switches locations on the external memory throughout sequence production.

Using the perceptron analysis we computed how the number of *output* gate neurons depends on task parameters SI Section 6.1.5), it has a term roughly proportional to the number of roles (Fig. 3d) and it scales as the square root of the number of symbol each role can be filled with (Fig. 3e). With the theory, we can also show that splitting neural representations for each role into segregated networks, as observed in prefrontal cortex (18), is more efficient than storing distributed representations of all role-filler pairs in a single attractor network with multi-item working memory (Fig. 3f, SI Section 6.1.5).

Our automaton interpretation for this neural architecture is the following: the NLSC encoding the abstract sequence is a FSA that implements a reading head which couples the relevant cell of the external memory with a machine printing the currently represented symbol (Fig. 3g). From this perspective PFC circuits of the monkeys implement an external memory accessed by e.g. lower motor centers. It would be interesting to see whether neurons implementing the head NLSC for the abstract sequence XYX, or *output* gate neurons with a mixed selectivity to roles and fillers, could be identified in cortical networks of animals performing such a task.

### 4 Assembling NLSC for behavioral programs

#### Neural Turing machines

Here we show how this ability to move on the external memory (Fig. 3g) can be combined with the ability to write on it (Fig. 2e). The external memory is now composed of multiple NLSC (Fig. 4a shows a full NLSC with both populations of transition and output gate neurons), each implementing a memory cell (Fig. 4b green). A NLSC plays the role of a FSA, or machine, reading and writing on the external memory (Fig. 4b blue). A NLSC plays the role of the head, deciding from which cell the machine is reading and writing (Fig. 4b purple). Thus, combining NLSC allows to assemble Turing machines (Fig. 4c, SI Section 6.5). Turing machines are fundamental to computer science. They consist of simple automata, which manipulate symbols according to sets of rules. Yet, they are computationally universal, meaning that such simple machines can implement any definable procedure, algorithm, or program. For systems neuroscience, it implies that NLSC-based architectures can support any pre-defined behavioral program performed by an animal in an experiment. This statement implicitly assumes that new memory cells can be added on demand during computation, so as to effectively obtain an infinite external memory (see Discussion). Such a construction allows us to define the notion of behavioral program. As input to the program, the external memory is loaded by setting the dictionary of each of its NLSC into persistent states representing cells’ symbols. Transitions between machine, head and cells states correspond to computation unfolding over time. The output of the program is the final state of the tape or of the machine.

**Fig. 4:**
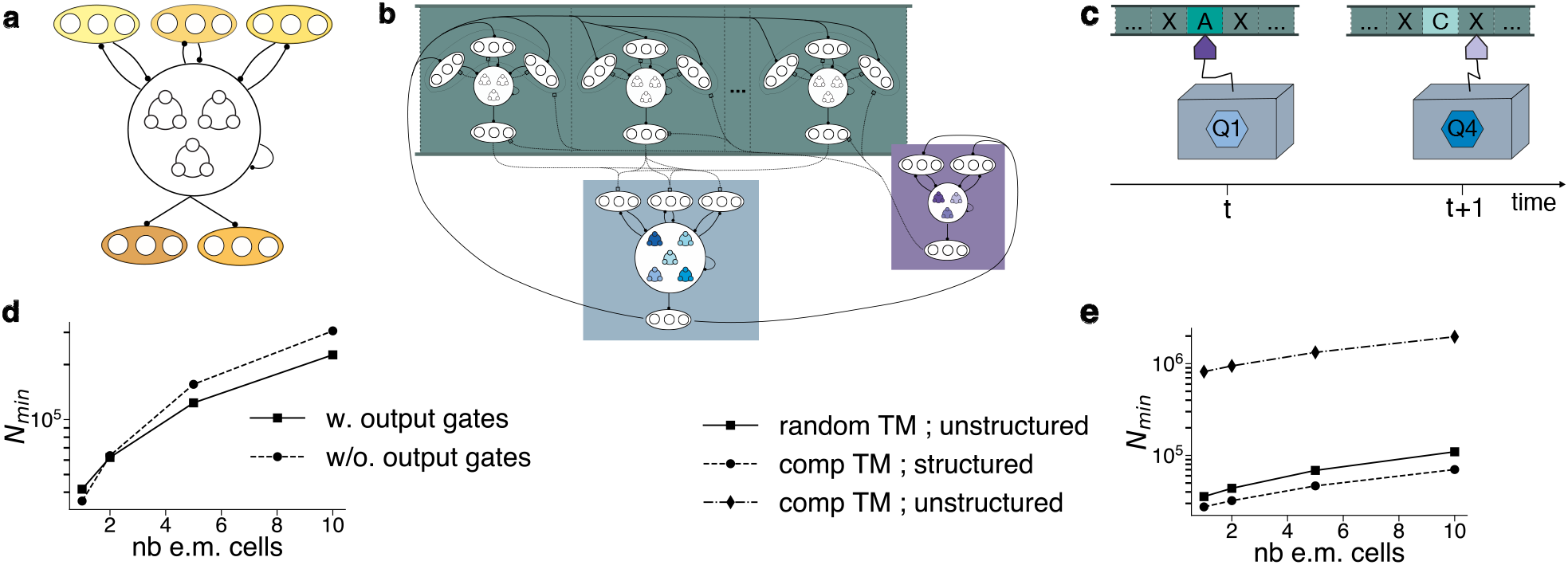
Assembling NLSC for the implementation of behavioral programs. **a**, NLSC with both populations of transition gate neurons (top, yellow) implementing multiple permutations of dictionary states, and populations of output gate neurons (bottom, orange) allowing to route information transmission to downstream circuits. **b**, Neural Turing machine, **c** mapped onto its conventional automata representation. **d**, N_min_ as a function of the number of cells composing the external memory, for NLSC-based Turing machines with and without the output gate neurons. **e**, N_min_ as a function of the number of cells composing the external memory, for NLSC-based Turing machines implementing random transitions with non-compositional neural representations (squares), structured transitions with compositional neural representations (circles), and random transitions with compositional neural representations (diamonds).

#### Assembling NLSC for a specific behavioral program

To illustrate how to assemble NLSC to build such an architecture, with both moving and writing on an external memory, we considered a foraging task with a planning component. An agent is navigating on a map of 10 spatial locations linked through 3 different paths (Fig. 5a, blue arrows). A value, representing e.g. food quantity is associated to each location, and the goal of the agent is to decrease its hunger level. When assembling NLSC we programmed the agent to implement a specific algorithm described by the pseudo-code in SI Section 6.3. This algorithm is illustrated in Fig. 5b-c. From each starting location (e.g. *X*_9_), the agent plans each of the three paths and, using a memory of the food values associated to each location, records the distance to the next food location (Fig. 5b illustrates planning for path 2). It then decides to move along the path with the shortest distance to food, travels and consumes food, decreases its hunger level, and start the planning process again if still hungry (Fig. 5b).

**Fig. 5:**
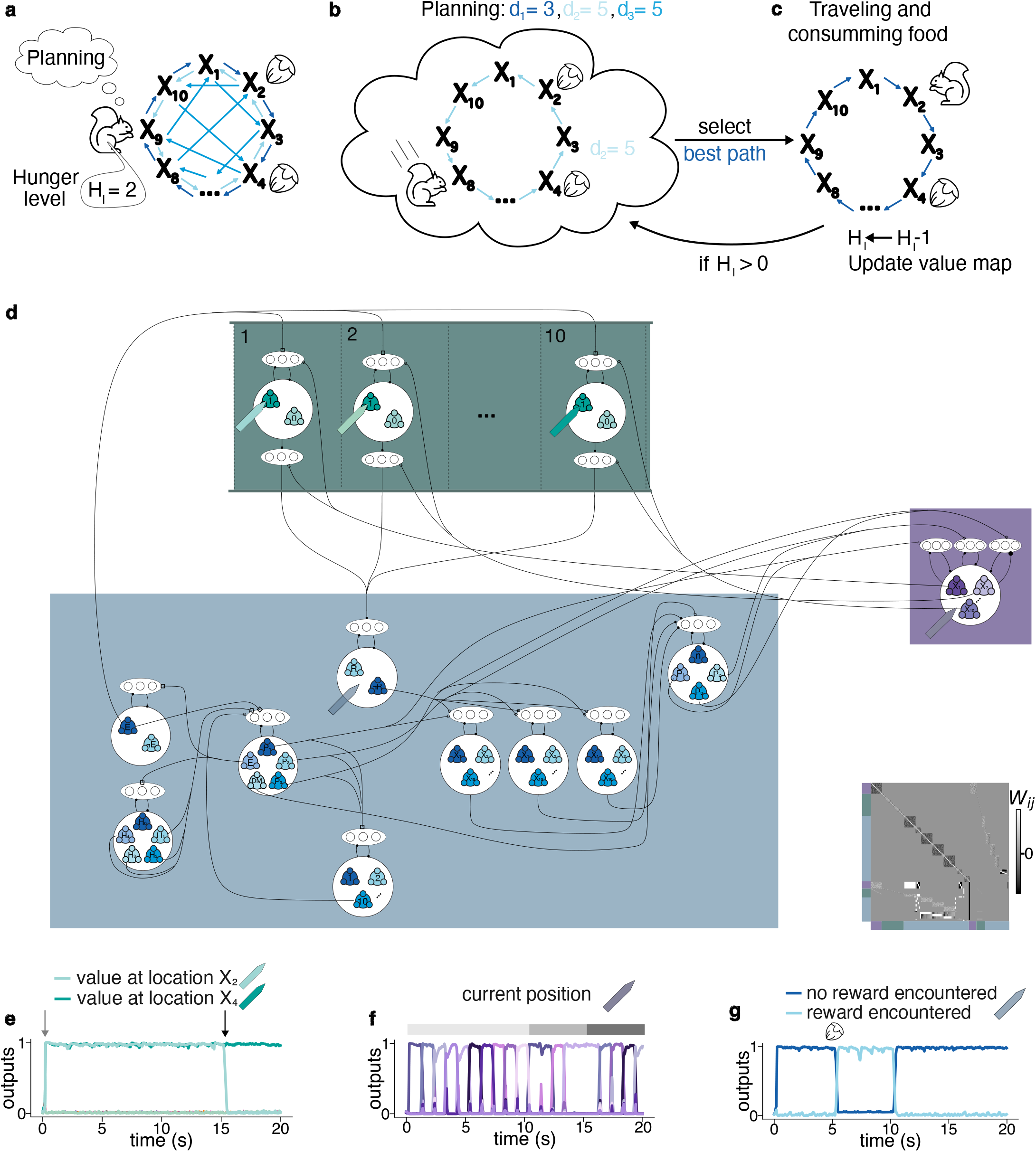
Solving a foraging task with a neural Turing machine. **a**, Environment on which an agent evolves to satisfy a hunger level, 10 locations are linked by three paths (blue shaded arrows), food can be present at each location. **b**, Example of a planing step of the behavioral program, whereby the agent mentally travels on the light-blue path 2 to estimate the distance from its current location to the next location with a food item. **c**, After selecting the path with shortest distance from food, the agent travels, consumes food, updates its hunger level as well as its mental representation of where food is available. If still hungry it goes back to planing a route towards a new food location. **d**, Assemblage of NLSC to solve the task. **e**, Dictionary activations for the 10 NLSC of the external memory circuits (green electrodes in (d), readouts that signal 0 value are not shown): at the time of initialization (light arrow) food items are present at two locations. Once a food item is consumed the external memory cell coding value at this location is updated (dark arrow). **f**, Cell assembly activations for the head circuit (purple electrode) while planing path 3 (light gray), traveling to consume food (middle-light gray) and start planing again (dark gray). **g**, Cell assembly activations for the machine (blue electrode) updating its state based on the currently read cell content, the particular NLSC we show keeps a memory that food has been encountered in planing a path.

The final neural architecture is represented in Fig. 5d and its relationship with the pseudo-code is described in details in Method 5.3.5 and SI Section 6.4. We now describe cell assembly activations in the different sub-circuits while running the behavioral program (cartooned recording electrodes in Fig. 5d). The variables storing the food value associated to each spatial location are represented in the external memory using 10 NLSC with a 2-states dictionary (value 0 or 1). At the beginning of the task, the external memory is loaded with values, mimicking an agent recollecting memories of where food has been cached (Fig. 5e, light arrow, two locations with food). When, at some point during the behavioral program, food is consumed at a location X, the associated value is decreased by switching the dictionary state of the Xth NLSC of the external memory (Fig. 5e, dark arrow). In order to navigate on the 10 spatial locations, either during the planing components or while travelling effectively towards food consumption, the neural architecture is equipped with a head NLSC with a 10-states dictionary (one per location) and three populations of gate neurons, each programing one of the path. Fig. 5f shows cell assembly activations in the head circuit for planing path 3 (light gray), traveling to consume food (medium gray) and planing again on path 1 (dark gray). Each cell assembly activation in this circuit enables communication between one of the external memory cell and the machine circuit (connections from purple to green circuits Fig. 5d). The machine circuit is composed of multiple NLSC that coordinate the running of the behavioral algorithm (Method 5.3.5) or that hold variables in memory to program transitions. Fig. 5g shows the activity in such a NLSC, it encodes whether food has been encountered while planing a path, allowing the rest of the machine circuit to evaluate distance from food on each path (Method 5.3.5, SI Section 6.4).

This illustrates how states of a head circuit (Fig. 5d purple NLSC), couple a machine circuits (Fig. 5d blue NLSC) with different external memory circuits (Fig. 5d green NLSC). The posterior parietal cortex (43) or dorsolateral prefrontal cortex (44) could implement the NLSC of the head circuit representing and updating the (mental) position of the animal, while the external memory representing values for each position could be implemented by NLSC in the orbito-frontal cortex (45), whereby individual neurons would encode conjunctions of a specific position with the value associated with this position.

#### Capacity of neural Turing machines

In practice, as in any machine, the cells of the external memory are in finite number. The number of neurons *N*_*min*_ required to build a defined NLSC-based Turing machine with a finite and fixed number of memory cells can be computed (SI Section 6.1.6, SI Fig. 6e). Below we describe how *N*_*min*_ depends on the various parameters of a statistical ensemble of programs with randomly drawn transitions between states. It depends: for the external memory, on its number of cells and the number of symbols supported on each cell; for the machine, on the number of machine states; for the head system, on the number of cells. The size and number of gate populations depend on the number of dictionary transitions that should be coordinated among external memory, machine and head NLSC. In Fig. 4d, we show *N*_*min*_ for a random Turing machine with *P* = 1000 machine states and *P*_*e*.*m*._ = 100 external memory symbols (SI Section 6.1.6). Extending the tape from 1 to 10 cells increases *N*_*min*_ from 42, 000 to 200, 000 (full line). The dashed lines shows that, as long as there is more than one external memory cell interacting with the machine circuit, it is beneficial to modulate communications between NLSC through output gate neurons (bottom dark-orange neurons in Fig. 4a), rather than duplicating gate neurons of the machine NLSC.

In this modeling, machine states are carried by a single NLSC with *P* states {*X*^*µ*^} _*µ*=1,…,*P*_. This is not the case for the neural architecture we proposed for the foraging task, which has a machine circuit composed of multiple NLSC (Fig. 5d, blue), such that machine states are indexed not by P symbols but by tuplets of symbols (*path, hunger level, distance counter*…): representations are compositional, or factored. The dashed line in Fig. 4e shows that using compositional representations 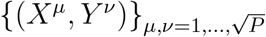 supported by 2 NLSC allows to reduce *N*_*min*_ from 110, 000 (full line, non-compositional representations) to 70, 000 (dashed-line, compositional representations) for implementing a program with a structured set of transitions between dictionary states (SI Section 6.1.6). In general compositional representations are not beneficial for unstructured sets of transitions (Fig. 4e, dashed-point line) since the number of gate neurons increases faster than the number of dictionary neurons decreases.

## Discussion

We put forward artificial neural networks performing a set of tasks with primitive temporal structures that have been proposed to underly behavior (23). This lead us to highlight the Neural Latching Switch Circuit (NLSC) as an elementary circuit (Fig. 4a) that can be combined to assemble neural architectures suited for all these tasks.

### Relationship between neural and symbolic computations

To characterize these neural architectures we leveraged neural network theory tools developed for the study of attractor neural networks (SI Discussion 6.8.1). Attractor neural networks (14) are described as dynamical systems in which trajectories of neural activity converge towards fixed-points, a feature that has been proposed to support various computations such as working-memory maintenance, stimulus denoising or temporal integration (13). In order to provide sensible definitions for temporal structures and for the computational power of our neural architectures, we resorted to computer science and interpreted attractor states as automata states (24).

In this framework, NLSC can act as the machine, the external memory, or the head of a Turing machine (Fig. 4b,c), showing it can be combined as a building-block to implement any behavioral program (SI Discussion 6.8.2). From this point of view, neural computations are instantiated by the coordinated transitions between attractor-states of multiple NLSC. This extends the framework of computation through neural dynamics (14; 46) and establishes a bridge between neural and symbolic computations, whereby neural network operations are conceptualized as symbol manipulations. We computed how the minimal number of neurons of an architecture depends on its computational power (e.g. number of external memory cells, machine states, head moves). This has allowed us to compare various architectures and to show that optimized NLSC-based architectures, which use distributed and sparse neural patterns (47; 48), are efficient, for instance when comparing our finite-state automaton implementation with the classic one proposed by McCulloch and Pitts (25; 24).

Recent works formulated a similar relationship between cell assembly activations and symbol manipulations (49; 50). In these implementations, cell assembly activations are equated with both dictionary and gate neural patterns of the machine circuit, while cell assemblies for the external memory cells and head circuit are not coupled to gate neurons. By contrast every element of our neural Turing machines is embodied in a NLSC, with cell assembly activations in dictionaries interpreted as machine states, symbols represented on each memory cell, or a cell address in the head circuit. Also, the functioning of the external memory is different, in particular it relies on fast synaptic plasticity for writing new symbols on a cell, while here, writing a new symbol corresponds to switching a dictionary to a new state, thanks to activations of gate neurons.

The approach exposed here and in (50) can be contrasted with the neural engineering framework (51), which also put forward a recipe to build neural networks suited for any behavior. It relies on the generic fact that neural networks are universal function approximators and can implement any dynamical system. Here, by going through computer science and automata, we propose a more constructive approach, exposing how an elementary neural circuit embodies minimal ingredients underlying computation. Furthermore, we deployed statistical physics tools to show how network sizes and structures relate to the specific computations to be implemented, allowing, in principle, quantitative comparisons with other theories or with experimental characterizations of brain circuits.

### Relationship with brain circuits

The concept of computation has proven valuable for understanding the function of various biological entities (52) and may as well offer insights into the workings of brain circuits. In (24) we have discussed how NLSC can be mapped onto the head-direction system described in the central complex of the fly’s brain (53; 26), with neurons in the ellipsoid body, selective to head-direction, playing the role of dictionary neurons, and neurons in the protocerebral bridge, selective to conjunction of head-direction and angular velocity, playing the role of gate neurons. In line with previous modeling studies (54; 21), we have also discussed thalamic neurons as a potential substrate for gate neurons. However we noted that the number of thalamic neurons is low compared to the number of cortical neurons, making cortico-thalamic circuits not well suited for the long-term storage of the procedural memories of NLSC. Another possibility is that cortical columns (55) can implement NLSC, with e.g. L2/3 neurons implementing transition gate neurons and L5/6 neurons implementing output gate neurons (Fig. 4a yellow and dark-orange). L2/3 neurons of primary motor cortex of rodents have been shown to mix representations of one of two task contexts with current motor kinematics (56), and could serve as transition gate neurons, directing sequences of neural representations driving the behavior associated with each context. In architectures implementing Turing machines or producing sequences with an algebraic pattern structure, output gate neurons under the control of the head circuit, are used to efficiently set-up communication channels between NLSC (Fig. 4d). It is tempting to identify these gate neurons with L5/6 neurons that have been proposed to serve as outputs of the cortical column (57; 58). Gain modulation of L2/3 or L5 neurons via long-range cortical inputs, implemented either through local micro-circuitry (59) or dendritic (60) non-linearities, could mediate interactions between NLSC. In our neural architectures, transition gate neurons are exclusively driving their target dictionary neurons. By contrast, output gate neurons can both drive or modulate their target, depending on whether it targets a NLSC with higher or lower number of dictionary states. It could be characterized experimentally whether L2/3 or L5 neurons exhibit different patterns of driving and modulatory effects on their target neurons.

Throughout the manuscript we have shown how interactions within large-scale brain circuits can be understood as interactions between NLSC. Interactions between pairs of areas in motor-cortex (39; 38) can be seen as implementing motor procedures with long-distance dependencies or organized into chunks (61) (Fig. 1b,Fig. 3b), with a higher-level area (green circuits) evolving on a slow timescale and controlling the paths taken by neural activity in a lower-level area (blue circuit) on a faster timescale. We also expect interactions between triplets of areas, implementing the head, machine and external memory circuits (Fig. 3b,Fig. 4b,Fig. 5d). For instance, in a saccade task with an algebraic pattern structure, neurons in prefrontal cortex can be seen as implementing cells of an external memory (green circuits), holding representations of multiple target locations during the maintenance epoch of the task (18). During the saccade production epoch of the task, each of this cell circuits could be read in succession by a machine circuit (blue circuit), such as the frontal eye-field known to encode the target location of saccades. This succession of couplings between one of the external memory cell and the machine circuit is expected to be mediated by a third head circuit (purple circuit), located in another brain area to be identified. Such triplets of interacting NLSC is also involved in the foraging task we devised to illustrate the implementation of behavioral programs. Based on known neural selectivity, we speculated that in this case posterior parietal cortex or dorsolateral prefrontal cortex could be seen as a head circuit, and that orbito-frontal circuits could be seen as an external memory, with neurons encoding conjunctions of location and value. Further work is required to reach more precise descriptions of these systems and see how well they map onto interacting NLSC, and e.g. see whether dictionary and gate neurons can be assigned to specific cell classes or not.

The quantitative theory we developed allows to describe the anatomical and physiological properties of efficient neural architectures. For a given behavior, a confrontation with experimental data can be devised. This should aim to characterize the selectivity of recorded neurons to task variables (28), describe whether neurons are driven or modulated by their various inputs (62; 59), characterize the coding level of cells and their cell class dependence (63), but also the trajectories of population recordings during task execution (64), the connectivity statistics between cell classes (65; 66) or the population structure of circuits (67; 68). The examples of behaviors treated here are inspired by animal models currently used in systems neuroscience experiments. Of particular interest are context-dependent tasks (34; 36; 69), which are known to engage interactions between distinct cortical areas and thus seem well suited to confront cortical networks with our theory. As for the fly’s head-direction circuit, understanding whether and how mammalian brain circuits implement path integration on continuous attractors could also be interesting, although the current theory might have to be extended to give adequate descriptions (24).

### Functional role for neural oscillations

In most of the neural architectures we discussed, an external go cue acted as a clock signal to trigger transitions among neural states. In (24) we suggested that network oscillations (12) could serve as internally generated clocks, leading attractor-states to become metastable (22). This is less of a strong role for oscillations than in binding by synchrony (70; 71), where oscillatory modes are proposed to dynamically establish communication channels between neural circuits. In NLSC architectures, communication channels are pre-wired, with contextual representations in higher-level areas deciding which communication channel to activate, while oscillations pace dictionary state updates. Further studies could assess whether brain oscillations observed at various frequencies (12) reflect updating of neural network states, similar to what has been observed for gamma frequencies (72) (SI Discussion 6.8.3). The highly-structured network architectures we presented in this work can also be contrasted with the unstructured networks described in (73), which, poised at a critical point can be reconfigured with inputs to perform different tasks.

### Temporal structures of behavior reflecting computational abilities of neural circuits

In trying to find a neural substrate underlying high-level cognitive processes, we defined a set of temporal structures that reveals computational features of the underlying neural circuitry (Fig. 6). To do so, we took inspiration from (23) which proposed chunking and algebraic patterns as two important forms of temporal structures to model behavior. Interestingly, we found that behaviors associated with these temporal structures rely, respectively, on the computational ability (25) of neural circuits to write symbols and to move on cells of an external memory.

**Fig. 6:**
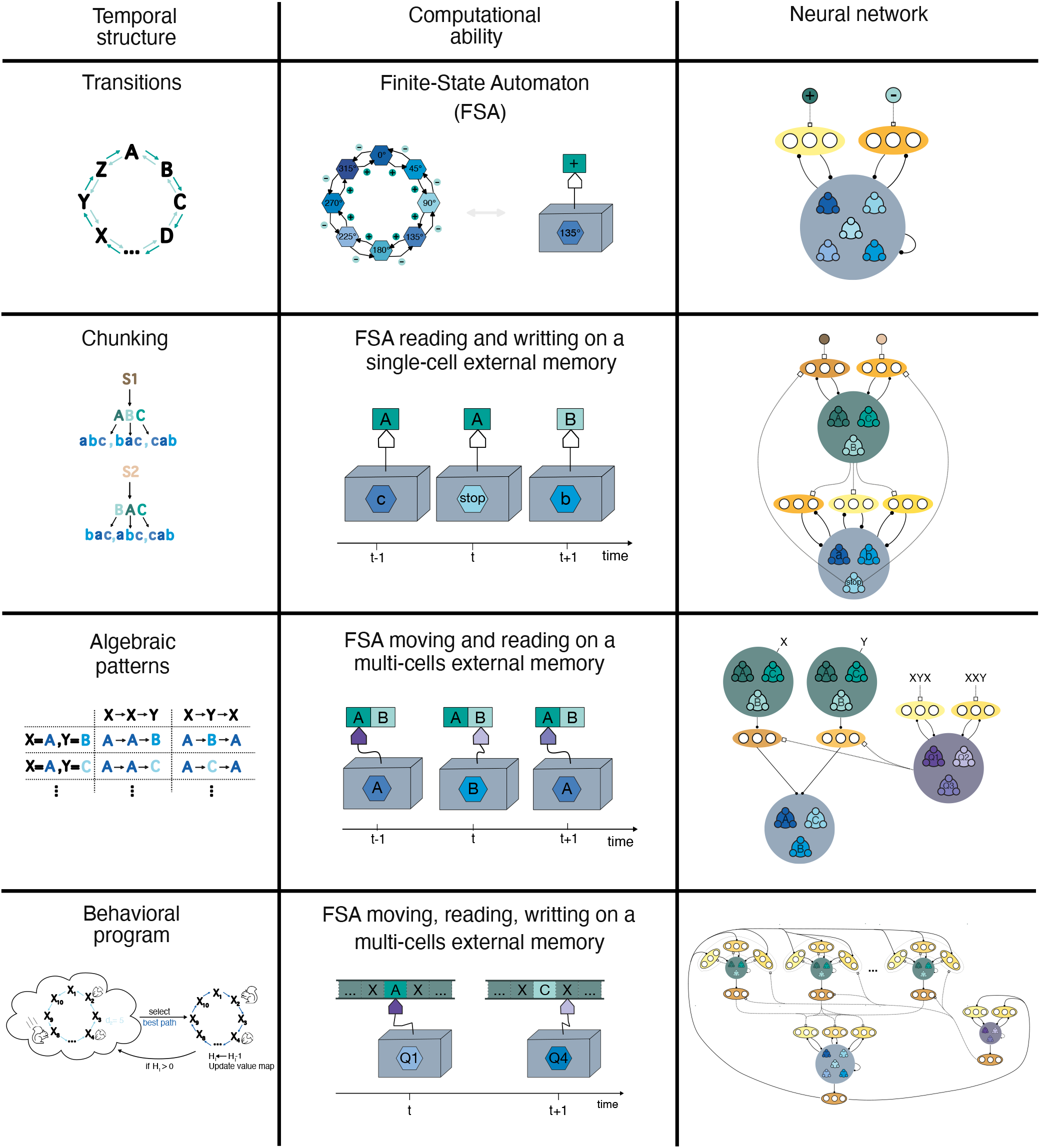
Relationships between temporal structures of behavior and computational abilities of neural networks. Left column: temporal structures used to describe behavior. Middle column: automata with various computational abilities. Right column: assemblage of Neural Latching Switch Circuits to physically implement automata. The different colors represent the different computational roles played by different NLSC.

Generative grammars, arranged in a hierarchy of complexity levels (SI Fig. 1d), and their correspondence with automata represents another attempt at linking behavior and computation. Generative grammar rules have been introduced to generate strings of symbols capturing properties of natural languages. However, the relevance of this hierarchy of temporal structures for modeling behavior has not been clearly exposed, for instance, natural language is not easily assigned to one of the complexity class of generative grammar rules ((8), Chapter 15.2). While neural networks such as NLSC can be well mapped onto finite-state automata, which correspond to regular grammars constituting the first level of the Chomsky’s hierarchy (SI Fig. 1a,b), it is more difficult to map neural network operations onto push-down automata, which are associated to the second level of the Chomsky’s hierarchy (SI Fig. 1c). This is to be contrasted with interactions between NLSC that can naturally be mapped onto automata abilities. Thus, considering physical implementations of automata into neural networks, sheds light on why generative grammar might not be well suited to model behavior.

Another temporal structure proposed in (23) involves the generation of nested-tree structures, accounting for the combinatorial expressivity of behaviors such as natural language. Here, guided by computational principles, we ended-up putting the focus on the notion of behavioral programs instead, which is also associated with a form of combinatorial expressivity (SI Section 6.7). Behavioral programs as conceptualized here are supported by external memories with a finite number of cells. As such, NLSC architectures are not able to emulate Turing machines with an infinite number of memory cells. However, brains have a finite number of neurons, and behavioral studies have shown that human working memory capacity (the number of memory cells maintaining an active representation) is rather limited. Future work will further detail how interacting NLSC can lead to the combinatorial power characteristic of cognition.

Previous studies have shown that imposing physical constraints on the underlying neural substrate can lead to quantitative predictions of behavioral measurements (74; 75; 76). Further work is required to investigate whether the computational account exposed here could be tested in similar ways, for example by predicting the behavioral circumstances under which compositional mental representations can be expected (77) (Fig. 4e), or by examining features of natural languages (SI Section 6.7).

## Acknowledgments

We are grateful to Arthur Leblois, Rémi Monasson and Gianluigi Mongillo for discussions and a careful reading of the manuscript.

## Data and materials availability

All codes, for training ANN, solving analytical equations and generating figures, will be made available on a public repository at the time of publication.

## 5 Methods

### 5.1 Artificial neural networks

We trained networks of rate units that are discretized version of the continuous time equations

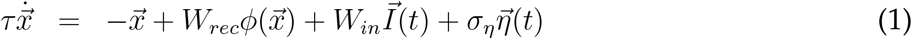

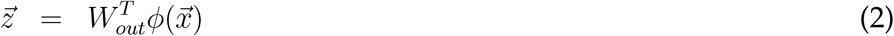

where 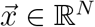 is the vector of currents to the *N* neurons composing the network. Currents are injected in the network by *N*_*in*_ input neurons 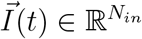 through *N*_*in*_ input weight vectors collected in the matrix *W*_*in*_ ∈ ℳ (*N*_*in*_, *N*). We divide inputs into two categories 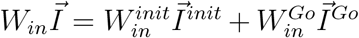, where 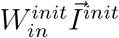 stands for input used to initialize networks into specific stable states and 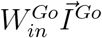 for inputs triggering transitions between stable states. Currents flow in the network through the recurrent matrix *W*_*rec*_ ∈ ℳ (*N, N*). These currents are transformed into firing rates through the activation function *ϕ*(.) which we take to be the rectified linear unit except for low-rank networks for which we used the hyperbolic tangent that leads to more interpretable mean-field equations. *N*_*out*_ output neurons 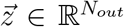 read linear projections of the firing rate vector 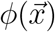 through *N*_*out*_ output weight vectors collected in the matrix *W*_*out*_ ∈ ℳ (*N, N*_*out*_). 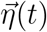 is a vector of independent white noises, with standard deviations *σ*_*η*_ = 0.05.

Training is performed using the Pytorch library by tuning specified connectivity parameters so as to minimize a loss 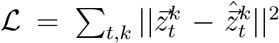, where *k* indexes trials in the input-output dataset and *t* indexes discretized time (we used a discretization time step of 20ms, for a time constant *τ* = 100ms). 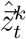 represents the desired output activations, while 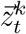 represents the actual output activations. Minimization is performed with stochastic gradient descent using the Adam optimizer with standard parameters. The choice of connectivity parameters used to minimize the loss depends on the specific training set-ups we used.

### 5.2 Training set-ups

#### 5.2.1 Training NLSC

Networks are segregated into *N* = *N*_*dic*_ + *N*_*gate*_ neurons through constraints on the connectivity (SI Fig. 2a). The *N*_*dic*_ neurons receive inputs from the initializing inputs 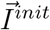 and are connected to the readouts, while the *N*_*gate*_ neurons receive inputs from the transition inputs 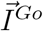 and do not connect to the readouts. The *N*_*dic*_ dictionary neurons are recurrently connected with each others, while the *N*_*gate*_ gate neurons are not, though gate neurons are bidirectionally coupled with dictionary neurons. Dictionary connectivity parameters 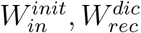 are trained on the dictionary task to implement *P* stable states (Method 5.3.1) while output connectivity *W*_*out*_ are fixed with readout neurons receiving from *P* non-overlapping subsets of dictionary neurons to ease interpretation of trained networks. Input connectivity to gate 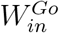 and gate/dictionary connectivities *W*^*dg*^, *W*^*gd*^ are trained to implement task-specific transitions between dictionary states. All initial connectivity parameters are drawn from Gaussian distributions. The time constant of gate neurons is decreased from 100 to 20ms to obtain sharper state transitions. Except when notified, *N*_*dic*_ is equal to *P ×* 41 and the number of gate neurons equals the number of dictionary neurons.

#### 5.2.2 Training unconstrained networks

Neurons are not segregated into dictionary and gate neurons, and we train an unconstrained recurrent matrix *W*_*rec*_ (SI Fig. 2b) to implement stable states and task-specific transitions. Input weights 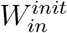 and output weights *W*_*out*_ are fixed to values obtained by independently training a network on the dictionary task, while transition inputs 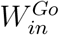 target all neurons with a fixed connectivity weight drawn from a centered normalized Gaussian distribution.

#### 5.2.3 Training low-rank networks

Recurrent matrices of rank-K are parametrized as 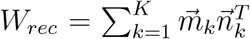 with 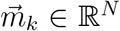. The rank K corresponds to the number of dictionary states, and connectivity vectors 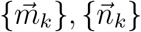 are initialized as Gaussian vectors structured into populations ((78), SI 6.2) to instantiate the relevant neural architectures. Each population is composed of 512 neurons. Values of the connectivity vectors are then fine-tuned to perform the task of interest by performing gradient descent on the loss parametrized with the 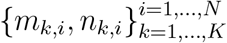 instead of the {*W*_*ij*_}_*i,j*=1,…,*N*_.

#### 5.2.4 Assembling NLSC

Here we describe how we obtained the networks in Fig. 2,3,5. The overall process consists in splitting the behaviors into individual transitions and then in training individual NLSC with appropriate representations and transitions, wiring different NLSC with each others with our understanding of population structure of gate neurons (24), and fine tuning the connectivity by performing gradient descent on longer and longer behavioral sequences (SI Fig. 2d).

In order to train individual NLSC we use the procedure of 5.2.1 with the addition that gate populations are segregated into sub-populations, one per dictionary states permutation. Two NLSC are then connected with each others by setting the connectivity from cell assemblies of the dictionaries to gate sub-populations. We do so by specifying the initial wiring and fine-tuning it by training the circuit on individual sub-tasks that contains coordinated transitions in the two NLSC. For networks assembled to perform the tasks of Fig. 5, we augmented the stability of dictionary states by adding a saturating non-linearity to the rectified linear activation function using *ϕ*(.) = *Hardtan*[0, 1](.), which allowed ANN to reliably perform the large number of transitions required for running the associated program. We also reduced the number of neurons per dictionary states from 41 to 20 to keep a relatively small network size. Programs and sequences are then produced thanks to an external go cue that triggers a programmed transition each time it is activated.

For each task for which we used this procedure, we provide, in the tasks descriptions below, more details about how NLSC are build and wired to each others.

### 5.3 Descriptions of individual tasks

In all tasks, networks are first trained on trials corresponding to individual transitions making up the behavior of interest. These trials have a fixation epochs of duration *T*_*fix*_ = 100ms, followed by a *cueing epoch* of duration *T*_*cue*_ = 100ms where the dictionary is set into its initial state through activation of input neurons 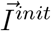. It is followed by a *maintenance epoch* of durations randomly drawn at each trial *T*_*main*._ ∈ [500, 2500]ms and a *go epoch* of duration *T*_*go*_ = 200ms during which the go signal is applied to the network 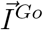, followed by a *response epoch* of duration *T*_*resp*_ = 1000ms. Networks are then retrained to perform the longer sequences in single trials in order to avoid dying out of stable states.

#### 5.3.1 Dictionary task

In order to build a dictionary of *P* states, we define an initializing input vector 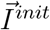, and a readout vector 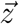 of size *P*. A single input neurons *j* ∈ [1, …, *P*] is activated at a value 1 during the *cueing epoch* if the trial starts with maintaining dictionary state *X*_*j*_. We set the target activations 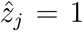 and 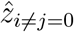. Here, and only for this task, we do not introduce additional transition inputs 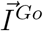, rather initializing input neurons 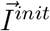 can also be activated during the *go epoch*. If input neuron *k* ∈ [1, …, *P*] is activated at 1 during the go epoch, we trained networks with target output activations 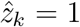 and 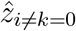 during the response epoch, while if the input neuron is activated at 0.5, 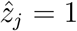 and 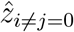 (SI Fig. 2c).

Networks trained on this task develop positive feedback loops among neurons belonging to the same cell assembly, defined as neurons projecting to the same readout neuron, as well as inhibition among neurons belonging to different cell assemblies. This connectivity structure leads to attractor dynamics making dictionary states robust stable states (24).

#### 5.3.2 Long-distance dependencies

In a first task we consider five symbols *A, A*^′^, *B, C, C*^′^ that have to be produced in two sequences *S*_1_ → *ABC* and *S*_2_ → *A*^′^*BC*^′^. The task structure is depicted in Fig. 1a. Activation of one of the initializing input neurons 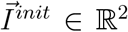 cues the network as to whether it should produce *A* or *A*^′^. Upon a first activation of the single go input neuron *I*^*Go*^, the network is asked to transit to state *B*, and then to *C* or *C*^′^ upon a second activation. In order to train on a task with long-distance-dependencies relying on more transitions (SI Fig. 3), we introduced a second task on a set of *P* = 20 states with initializing inputs 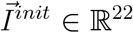 setting the network in one of the state and indicating at each trial one of two contexts by activating either input neuron 21 or 22 during the *cueing epoch*. Again the 20 states are thought of as being arranged on a ring, and depending on context, the network is asked, upon activation of a single go cue input, to transit in the clockwise or counter-clockwise direction. We first trained networks on individual transitions, and then on longer sequences up to 20 states.

A 5-states NLSC augmented with an external-memory trained on the first task has been presented in Fig. 1b,c. In order to obtain this network, we initialized the network with the NLSC, with random dictionary-gate connectivity, and added two populations of 41 neurons each preferentially activated by inputs activating either *A* or *A*^′^. The network is then trained to perform the task using Method 5.2.2. We showed how gate neurons activated in a context-dependent manner upon transiting out of state *B*. Manipulating these gate neurons allow to control transitions: in a trial *S*_1_, inhibiting the most active gate neurons in transitions *B* → *C* (yellow neurons in Fig. 1c, right) and exciting the most active neurons in *B* → *C*^′^ (orange neurons in Fig. 1c, right) triggers a transition from *B* to *C*^′^ instead of *C* (not shown).

For the second task, in figures SI Fig. 3e,f, we show the neural activity of a NLSC augmented with an external-memory while producing sequences of 20 states in either of the two contexts. SI Fig. 3b,c shows an analysis of gate patterns at transitions. Similarly to networks trained on the head-direction task (24), upon training, gate neurons segregate into two populations. With each population being preferentially inhibited by neural activation encoding memory maintenance of one of the two contexts. Inactivations during the *go epoch* confirmed that the observed population structure of gate patterns supports context-dependent transitions SI Fig. 3d.

We also trained unconstrained networks on this task (Method 5.2.2). In SI Fig. 3i,j we show the neural activity of such a network in the two contexts. From the rasters of neural activity, it is difficult to grasp how the memory of context is maintained throughout production of the sequence. SI Fig. 3k shows that the two neural states corresponding to the production of symbol *X* in the two different contexts are different, as opposed to NLSC networks SI Fig. 3g. The contextual memory is encoded in modulations of firing rates as observed in (27).

#### 5.3.3 Chunking

Here we explain how we built the neural architecture of Fig. 2b by assembling NLSC (Method 5.2.4). The full task is to produce two sequences *S*_1_ and *S*_2_. Each sequence is composed of *P*_2_ = 8 chunks *A, B, C, D, E, F, G, H* played in reverse order in *S*_1_ and *S*_2_. Each chunk corresponds to the production of a sequence of *P*_1_ = 20 states, starting with the same start and stop states. For instance chunk *A* is associated with the sequence of states *Start, a, b*, …, *r, Stop*, and other chunks are obtained by permuting symbols *a, b*, …, *r* without repeat (SI Fig. 5a,b top). A network with 20 states is trained on the dictionary task. *P*_2_ = 8 populations of gate neurons are associated to this dictionary, each being trained to implement one of the 8 sequences on the dictionary, giving us the low-level NLSC (Fig. 2b, blue). A second high-level NLSC (Fig. 2b, green) with 8 states in its dictionary and 2 gate populations is trained to implement the forward and backward transitions between chunks. A third dictionary with two states, or two sustained external inputs, encode sequence identity. These networks are then coupled with each others in the following way: each cell assembly associated with a chunk inhibits all gate populations of the low-level NLSC except one; low-level states, except the stop state, inhibit the two gate populations of the high-level NLSC; each of the two gate populations of the high-level NLSC is inhibited by one of the two states encoding sequence identity. The full network is then trained to perform each individual transition simultaneously in the two NLSC in response to activation of the single go cue input neuron *I*^*Go*^ that targets all gate populations. It is then retrained to produce longer sequences of length 20 in response to 19 successive activations of the go cue.

#### 5.3.4 Algebraic patterns

Here we explain how we assembled the neural architecture of Fig. 3b. The task is to produce all possible sequences of readout activations of the form XYX, where the roles X and Y can be filled by one out of 5 symbols, *A, B, C, D, E*. We first build a dictionary with 5 states that we duplicated to obtain the two memory slots. Each dictionary connects to a set of *output* gate neurons. We trained a head NLSC with a 3-states dictionary and a single gate population to produce the sequence *Q*_1_ → *Q*_2_ → *Q*_3_. We then initialize the connectivity from the head NLSC to the output gate neurons, such that each state *Q*_*k*_ is inhibiting the gates associated to the memory slot that should not be played. The full network is then retrained to perform all the possible 3 symbols sequences. In order to have this network perform sequences with another pattern (e.g. XXY), we simply add a new gate population to the head NLSC in order for it to produce a new sequence (e.g. *Q*_1_ → *Q*_3_ → *Q*_2_).

#### 5.3.5 Foraging task

An agent is foraging for food in an environment that consists of *P* = 10 locations linked through *nb*_*paths*_ = 3 different paths (Fig.5a). The working memory of the agent is initially loaded with food values associated to each location. It then mentally travels along the 3 paths and counts, for each path, the distance *K*_*path*_ from its current location to the nearest position with food available. It selects the path with shortest distance from food, travels along this path and consumes food to decrease its hunger level. It repeats these steps until the hunger level reaches 0. The task is summarized in the pseudo-code of SI Section 6.3.

In order to assemble NLSC implementing this program, we proceeded incrementally as described in Method 5.2.4, such that a go cue triggers relevant transitions inside the head, external memory and machine circuits. The final network is cartooned in Fig. 5d. Generically, variables of the pseudo-code are each implemented by a NLSC, with a number of dictionary states corresponding to the number of states the variable can be in. Interactions between variables throughout the behavioral program are set by connectivity weights from and to gate neurons. NLSC are also used to temporally scaffold interactions between variables (e.g. implementing **for** or **while** loops). We provide further details on the assemblage of these circuits in SI Section 6.4.

### 5.4 Mean-field theory for low-rank networks

In order to visualize neural activity as trajectories in state space for networks producing two sequences with long-distance dependencies (SI Fig. 4), we built low-rank networks with Gaussian vectors segregated into populations (78). Their dynamical behavior **(1)** can be reduced to interactions between cognitive variables, which characterize the location of neural activity along directions 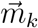:

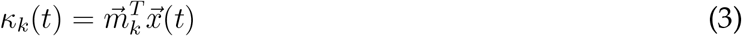

We have five cognitive variables *κ*_*A*_, *κ*_*A*_′, *κ*_*B*_, *κ*_*C*_, *κ*_*C*_′ associated with the symbols to be produced and two cognitive variables associated with the two hidden states maintaining the identity of the sequence in working memory *κ*_*hA*_, *κ*_*hA*_′. In particular the dynamics of *κ*_*C*_ writes

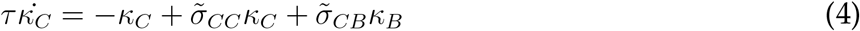

where 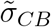 is modulated from 0 to 1 by the go cue only if (*κ*_*hA*_ = 1, *κ*_*hA*_′ = 0). There is a symmetric equation for *κ*_*C*_′, and other cognitive variables evolve as described in (24) Eq. **(4)**. Flow fields represent the time evolution of cognitive variables 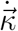 obtained from e.g. equation **(4)**, each panel is obtained using the *streamplot()* function of *matplotlib* on 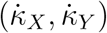.

### 5.5 Capacity of neural architectures

Our results on the minimal number of neurons, *N*_*min*_, required to implement NLSC-based architectures build on previous results obtained for perceptrons with binary neurons and binary synaptic weights (29; 30; 79; 31). We consider *N*_*out*_ perceptrons with *N*_*in*_ input neurons. The *P*_*a*_ input patterns 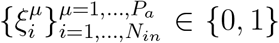 are drawn randomly with the constraint 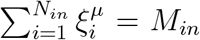 and output activations 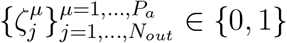 are drawn randomly.

We present calculations for computing the probability that the synaptic connectivity matrix 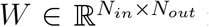 is not overloaded by the stored associations, i.e. that the output neurons correctly classify the *P*_*a*_ input patterns: 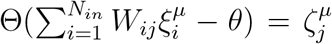, for all *µ* = 1, …, *P*_*a*_ and *j* = 1, …, *N*_*out*_, with Θ(.) the Heaviside function and 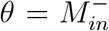 the activation threshold. The matrix of binary synaptic weights is assumed to undergo Hebbian learning while the *P*_*a*_ correct input and output activations are imposed onto the network: 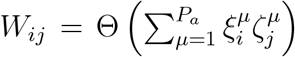. We note *g* the fraction of synapses activated after presenting *P*_*a*_ associations. It is a crucial quantity for our analysis as it controls the amount of interferences between associations and indicates for which value of *P*_*a*_ the learning rule is not able to correctly store the associations anymore. It can be estimated as

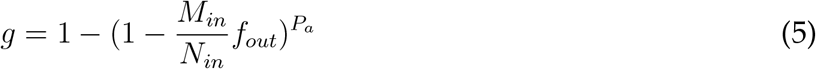

The probability that *P*_*a*_ associations in *N*_*out*_ perceptrons are learnt without errors can be expressed as (SI Section 6.1.1):

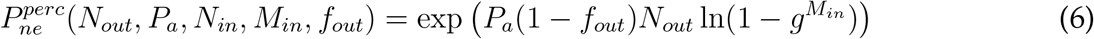

The neural architectures we built can be decomposed into multiple independent such perceptrons ((24) Fig.3a, SI Fig.7a-c). The probability that all the associations of an architecture are learnt without errors, 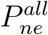, can thus be estimated by a product of the functions **(6)** estimated with the appropriate input variables. In order to estimate *N*_*min*_ for a given architecture and given number of associations, we performed grid-search over the number of neurons and coding properties *N*_*in*_, *M*_*in*_, *f*_*out*_ and look for sets of parameters that minimize the total number of neurons while keeping the probability of no error above a threshold 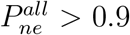. For instance, an attractor neural network of *N* neurons can be seen as *N* perceptrons. For *M* active neurons per pattern and *P* patterns, we compute 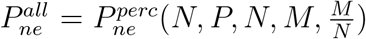 and perform grid-search on parameters *N* and *M* to find the values that minimize *N* keeping 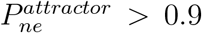. In SI Section 6.1, we detail this process for each of the neural architectures considered in the main text.

## 6 Supplementary

### 6.1 Capacity calculations

#### 6.1.1 Capacity for perceptrons

We detail how to obtain the probability **(6)** that the *P*_*a*_ associations are perfectly retrieved for *N*_*out*_ perceptrons with input size *N*_*in*_, input coding level 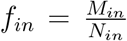 and output coding level *f*_*out*_. We do so following (31). To simplify mathematical expressions we have considered the case where the number of active neurons in the input patterns is fixed and does not fluctuate from pattern to pattern: 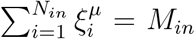 for all *µ*’s. With this constraint, the activation threshold of the output neurons can be taken as 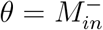, and, given the learning rule leading to

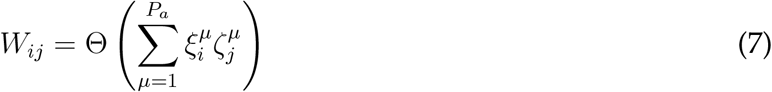

we are guaranteed that no errors will be produced when retrieving the *P*_*a*_*f*_*out*_ associations with 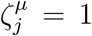. Errors can then only occur for associations with 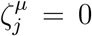. They occur when the output neuron receives an input *h* = *M*_*in*_, i.e. when all the synapses between the output neuron and active input neurons in 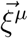 have been potentiated by one of the other *P*_*a*_ − 1 associations. The probability of such an event can be simply estimated by

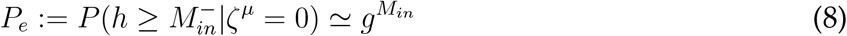

with *g* taken from **(5)**. The probability that no errors are produced in all the *P*_*a*_ associations and *N*_*out*_ perceptrons is then given by

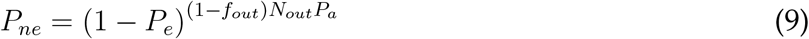

for sparse enough coding levels 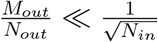 correlations between two synapses 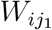 and 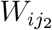 can be ignored (80) and the probability of error is well estimated by **(8)**.

#### 6.1.2 Storage capacity for single NLSC

Here we recap on the calculations presented in (24), where we showed how to compute the computational capacity of NLSC. An attractor neural network with *N* neurons can be decomposed into *N* single-output perceptrons with input size *N*, leading to a probability of no error for storing *P* patterns

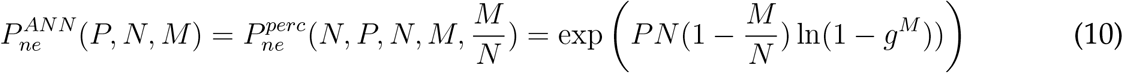

We described two architectures for a NLSC with *P* dictionary states, *P*_*c*_ contextual inputs, *P*_*c*_ × *ϵ P* transitions, *N*_*dic*_*/N*_*gate*_ dictionary/gate neurons and patterns of activity characterized with *M*_*dic*_*/M*_*gate*_ active dictionary/gate neurons. In the first architecture, dictionary neurons drive patterns of activity onto gate neurons, while contextual inputs modulate *P*_*c*_ populations of gate neurons. This architecture is efficient in the regime *P*_*c*_ ≪ *P*. In this case, the probability of no error for all the *P* stable states and all the *P*_*c*_ *× ϵP* transitions can be written as

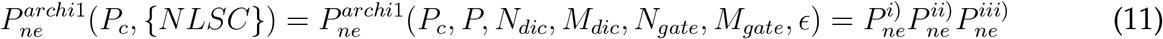

with

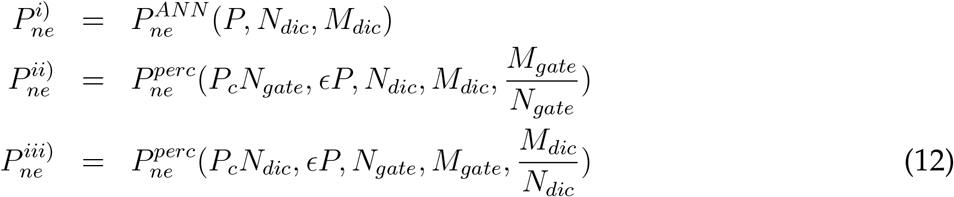

where {*NLSC*} = {*P, N*_*dic*_, *M*_*dic*_, *N*_*gate*_, *M*_*gate*_, *ϵ*} contains all the parameters defining structural and coding properties of the NLSC.

In the second architecture, contextual inputs drive patterns *P*_*c*_ patterns of activity onto each population of gate neurons, while dictionary neurons modulating *P* populations of gate neurons. This architecture is efficient in the regime *P*_*c*_ ≫ *P*. In this case, the probability of no error for all the *P* stable states and all the *P*_*c*_ *× ϵP* transitions can be written as

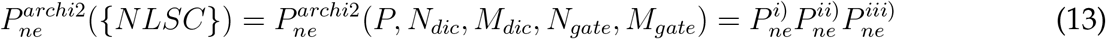

with

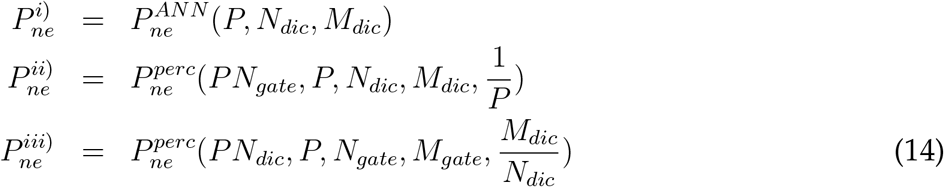

#### 6.1.3 Capacity of architectures for sequences with long-distance-dependencies

We consider *P*_2_ contextual signals that, instead of being carried by non-distributed representations, are distributed over the *N*_2_ neurons of an external memory with a number of activated neurons per patterns *M*_2_ (SI Fig.6a). We ask the patterns of neural activity in the external memory to be stable, i.e. the external memory is an attractor network. We now label *N*_1_ and *N*_1*g*_ the number of neurons in the dictionary and in each of the gate population of the NLSC, *M*_1_ and *M*_1*g*_ are the number of active neurons in dictionary and gate patterns, *P*_1_ is the number of dictionary states: {*NLSC*_1_} = {*P*_1_, *N*_1_, *M*_1_, *N*_1*g*_, *M*_1*g*_, *ϵ*_1_}. Compared to the section above, we also have to take into account interferences between connectivity patterns from the external memory to the gates of the NLSC, for the regime *P*_2_ ≪ *P*_1_, the overall probability of having no-errors can be written as

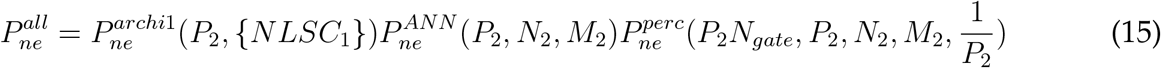

where the last two terms correspond to errors in imprinting *P*_2_ attractors in the external memory and errors in modulating the *P*_2_ populations of gate neurons (gate neurons have an activation threshold *θ* = *M*_1_ + *M*_2_). We have used *ϵ* introduced above, the fraction of transitions *T* = *ϵP*_1_ associated with each contextual signal.

For the case *P*_2_ ≫ *P*_1_, we have

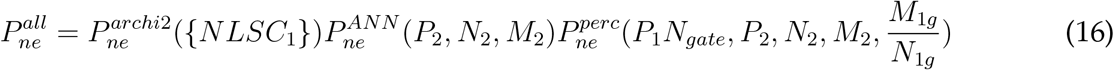

In the main text Fig.1e, we show *N*_*min*_ obtained from equations **(15)** (full line) and **(16)** (dashed line). We compared these values of *N*_*min*_ to those obtained by considering a neural architecture composed of *P*_2_ NLSC with *P*_1_ states in their dictionary (dotted-dashed line) for which

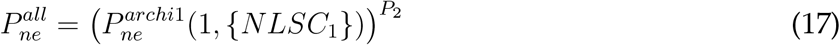

#### 6.1.4 Capacity of architectures for sequences organized into chunks

In the previous section, distinct states of the external memory provide distinct contextual inputs onto gate neurons to decide on transitions taking place in the lower-level NLSC. Instead of having only this uni-directional interaction, we would like the dictionary state of the lower-level NLSC to also trigger transitions in the external memory. To do so, we augment the external memory with gate neurons, giving us two interacting NLSC (SI Fig.6b). Each of the *NLSC*_*i*_, *i* ∈ {1, 2} has *P*_*i*_ dictionary states and is composed of *N*_*i*_ and *N*_*gi*_ dictionary and gate neurons. Each state in *NLSC*_*i*_ is associated with *T*_*j*_ = *ϵ*_*j*_*P*_*j*_ transitions in *NLSC*_*j*_. Depending on whether *P*_2_ ≪ *P*_1_ or *P*_2_ ≫ *P*_1_ it will be more efficient to have dictionary states of *NLSC*_2_ to modulate populations of gate neurons and *NLSC*_1_ to drive them or the converse, here we arbitrarily take *P*_2_ ≪ *P*_1_. We have

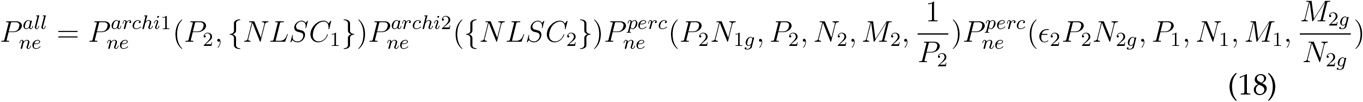

where the last two terms correspond to the connectivity from the dictionary of *NLSC*_1_ to the gates of *NLSC*_2_ and from the dictionary of *NLSC*_2_ to the gates of *NLSC*_1_.

In order to obtain the full curve in Fig. 2d, we consider a specific case where i) a single of *NLSC*_1_’s states, the stop state, triggers transitions in *NLSC*_2_ and ii) the ensemble of *ϵ*_1_*P*_1_ transitions in *NLSC*_1_ associated with a state in *NLSC*_2_ are closed (e.g. *A* → *O, O* → *Z, Z* → *E*, not *A* → *O, B* → *P, C* → *Q*). In order to point at the fact that this construction can be iterated hierarchically, we also require that the stop-state triggers closed transitions in *NLSC*_2_, and we take a total of *P*_3_*N*_2*g*_ gate neurons associated with *P*_3_ different sequences in *NLSC*_2_. For the figure we take *P*_1_ = 31, *P*_2_ = 10, 000, *ϵ*_1_*P*_1_ = 3 + 1 and for each of the *P*_3_ sequences, the stop state is associated with *T*_2_ = 100 transitions in *NLSC*_2_: the system we built stores *P*_3_ sentences each composed of 100 words (out of 10, 000) each composed of 3 phonemes (out of 30). We found *N*_*min*_ for this architecture by minimizing *N*_1_ + *P*_1_*N*_1*g*_ + *N*_2_ + *P*_3_*N*_2*g*_ under the constraint 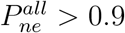, with

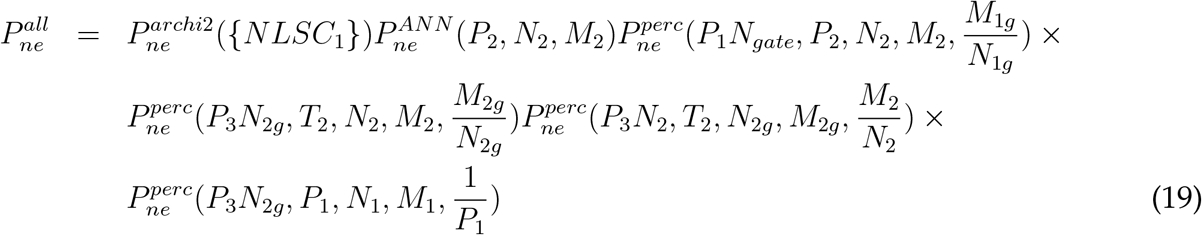

where the first three terms correspond to equation **(16)** for the articulatory apparatus, the next two terms correspond to connections in-between the dictionary and gates of *NLSC*_2_, and the last one to the connections from the stop state of *NLSC*_1_ to the gates of *NLSC*_2_.

We compared this implementation with one that does not make use of a stop state (Fig. 2d, dashed curve). *NLSC*_1_ has one dictionary state less. In order to program transitions in *NLSC*_2_ for relevant conjunctions of *NLSC*_1_ and *NLSC*_2_ states, gate neurons are segregated into *P*_1_ ≪ *P*_2_ populations and each of the *P*_1_ subpopulations will be involved with probability 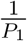 in triggering a transition in *NLSC*_2_ at the end of each word. The probability of no error associated with this architecture is obtained by introducing *P*_1_*P*_3_ populations of gate neurons in *NLSC*_2_:

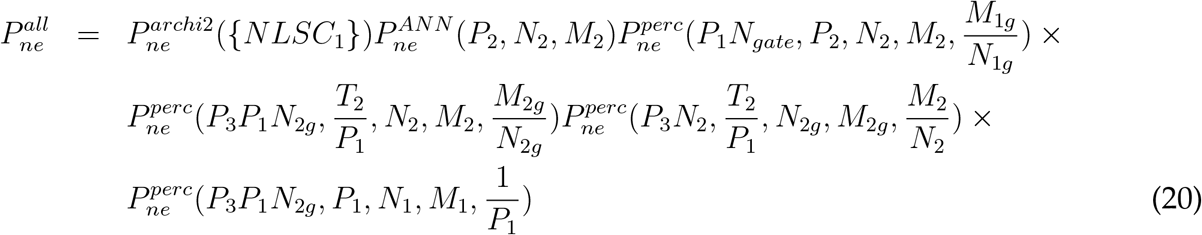

The scheme we presented for two hierarchical levels can be straightforwardly generalized to multiple levels by piling-up more than two NLSC and writing corresponding equations as in **(18)**.

#### 6.1.5 Capacity of architectures for algebraic patterns

We explain how we obtained the theoretical curves of Fig. 3d,e,f, that quantify the number of neurons used to produce sequences with a temporal structure characterized by algebraic patterns. We consider an external memory composed of *K* cells, each composed of a dictionary with *P* states, associated with *N*_*gate*_ *output* gates neurons driven by the *N*_*dic*_ dictionary neurons (SI Fig.6c). These *K* populations of gates then converge onto an output network of size *N*_*o*_ carrying neural representations used for production. In order for these representations to be manipulated by another downstream circuit we ask them to be stabilized by recurrent connectivity, i.e. the output neurons form an attractor network. The probabilities of no errors for this neural architecture is

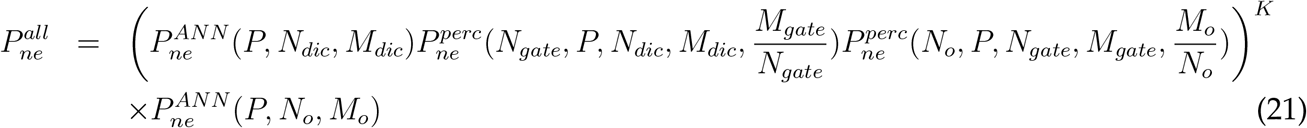

We compared this architecture with an alternative one that uses a single attractor neural network of size *N* with K-items working memory, instead of using K distinct single-item working memory attractor networks. In Fig. 3f, we report a lower bound on the minimal number of neurons that have to be used to maintain these representations. The network stores a total of *KP* patterns with *M* active neurons per pattern. We want to test the stability of a superposition of *K* patterns. As for the single item case, given the learning rule **(7)**, we are guarantee there is no error on foreground neurons. The probability of no error on background neurons is estimated as follows:

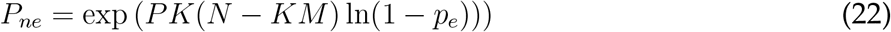

with *N* − *KM* the estimate of the number of background neurons, *PK* is the total number of pattern on which we impose there is no error. Note this number is smaller than the total number of possible patterns *P*^*K*^ on which the network can be in, i.e. we only evaluate the effect of increased probability of error on background neurons *p*_*e*_ due to increased activity from *M* to *KM*

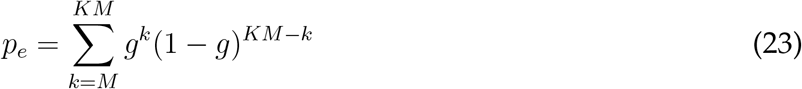

with

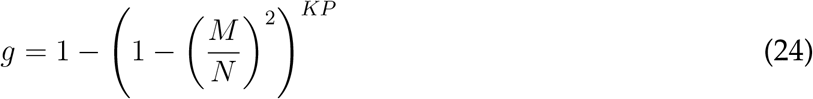

The number of gate neurons associated to the multi-item working memory is expected to be even higher than for the other architecture: the number of gate populations also scales as *K* given the need for gate representations to be modulated by the *K* states of the head circuit, and for each of these population, the associations are more difficult to implement given the increased amount of interferences in multi-item working memory representations.

#### 6.1.6 Capacity of neural Turing machines

The neural Turing machines we built are composed of three circuits (Fig. 4b):

An **external memory** circuit composed of *K* cells embodied in *K NLSC*_*e*.*m*._. We note *P*_2_ the number of symbols that can appear on each of the cells, *N*_2_ the number of dictionary neurons in each NLSC (*M*_2_ the associated number of active neurons composing a cell assembly), *N*_*g*2_ the number of *transition* gate neurons per sub-population used to implement transitions in each NLSC (coding level *M*_2*g*_) and *N*_2*g*_′ the number of *output* gate neurons used to transmit the currently supported state to the machine circuit (coding level *M*_2*g*_′).

A **machine** circuit (implementing a finite-state automaton) embodied by a single *NLSC*_*m*_. We note *P*_1_ the number of machine states, *N*_1_ the number of dictionary neurons in the NLSC (coding level *M*_1_), *N*_1*g*_ the number of transition gate neurons per subpopulation used to implement machine transitions (coding level *M*_1*g*_).

A **head** circuit embodied by an *NLSC*_*h*_ with a K-states dictionary. Gate neurons used to implement transitions between dictionary states. These gate neurons can be split into subpopulations of size *N*_*gh*_ to implement *P*_*h*_ distinct permutations of dictionary states, with a dictionary of size *N*_*h*_. For instance, a head circuit with two subpopulations, one associated with the +1 shift the other with the − 1 shift, would implement left/right head movements as for the canonical Turing machine. In order to limit the number of variables on which the optimization is performed to compute *N*_*min*_, we do not take the head-circuit into account. This is justified because we consider a parameter regime where the number of cells of the external memory *K* is small compared to the numbers of machine states *K* ≪ *P*_1_ and symbols of the external memory *K* ≪ *P*_2_.

Connections among these three circuits are as follows. The machine’s state conditions symbol transitions in a cell *k* of the external memory by connections from its dictionary to the transition gates of the *k*th NLSC. The state of the currently considered cell on the external memory is transmitted to the machine by connections from its output gates neurons to the transition gates of the machine. The head circuit routes communication between the machine and the *k*th NLSC of the external memory, this is done through connections from cell assembly *k* of its dictionary to transition and output gate neurons of *NLSC*_*k*_. Transitions in the head circuit are conditioned by machine states thanks to connections from the machine cell assemblies to the transition gates of the head circuit.

To estimate *N*_*min*_ for a Turing machine with *P*_2_ ≪ *P*_1_ (cf full line Fig. 4e), we implemented NLSC of the external memory with architecture 2 and the machine NLSC with architecture 1. Reusing previous calculations, the probability that there is no error in all the programmed transitions and stable states can be decomposed as

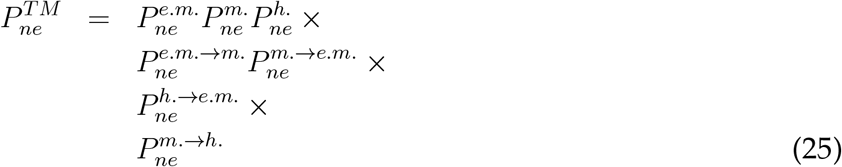

where the first line describes individual circuits, the second line describes connections between the external memory and machine circuits, the third line connections from the head to the external memory circuits, the last line the connections from the machine to the head circuit. Individual terms can be written as

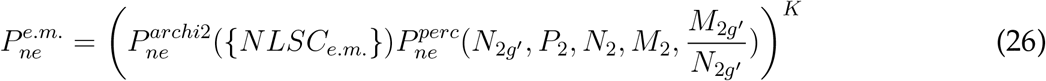

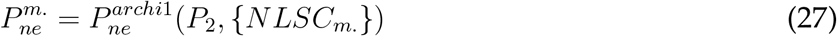

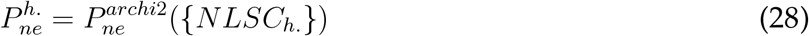

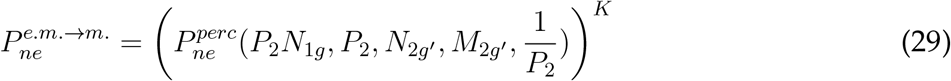

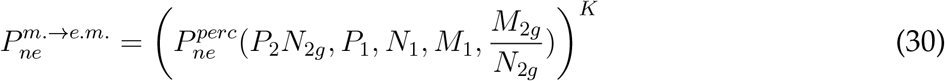

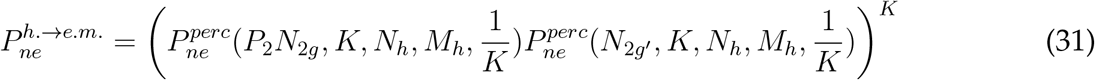

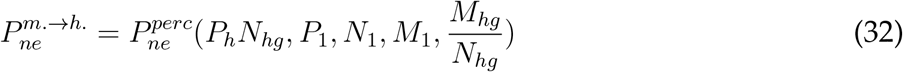

We used these equations to estimate how the number of neurons varies when varying various parameters describing the computational ability of the machine. The top-left panel in SI Fig.6e shows how the numbers of dictionary and gate neurons vary with the number of machine states, it is obtained for *K* = 5 external memory cells *P*_2_ = 100 symbols per cell. We optimized only over *N*_1_, *M*_1_, *N*_1*g*_, *M*_1*g*_, with fixed parameters {*NLSC*_*e*.*m*._}. The top-right panel shows variations of the number of neurons in the external memory with the number of symbols *P*_2_ for *K* = 5 and *P*_1_ = 1, 000. The bottom-left panel shows variation in numbers of neurons in the head circuit when the number of external memory cells *K* is varied, for a machine circuit with *P*_1_ = 1, 000. The bottom-right panel shows variation in the number of neurons in both the machine and external memory as a function of the fraction of transitions *ϵ*_*m*._ = *ϵ*_*e*.*m*._ = *ϵ*. As previously introduced, *ϵ*_*m*._ is the fraction of machine transitions associated to each symbol (at most there are *P*_1_ transitions per symbol) and *ϵ*_*e*.*m*._ is the fraction of external memory transitions associated to each machine state (at most there are *P*_2_ transitions per machine state).

In the main text we highlight that splitting the machine into multiple NLSC can be beneficial when the transitions to be programmed exhibit a form of structure. In order to obtain the dashed curves in Fig. 4e, we considered a machine circuit split into two NLSC with 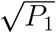 states each, for a total of *P*_1_ machine states. To obtain the red curve, corresponding to random transitions, gate neurons implementing transitions in the two machine NLSC and the external memory NLSC are split into 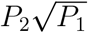 subpopulations, each positively gain modulated upon conjunction of a specific cell symbol and a specific state of the other machine NLSC. *N*_*min*_ is obtained by minimizing 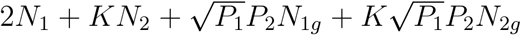 under the constraint 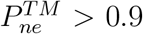 replacing equations **(26), (27)** of **(25)** with

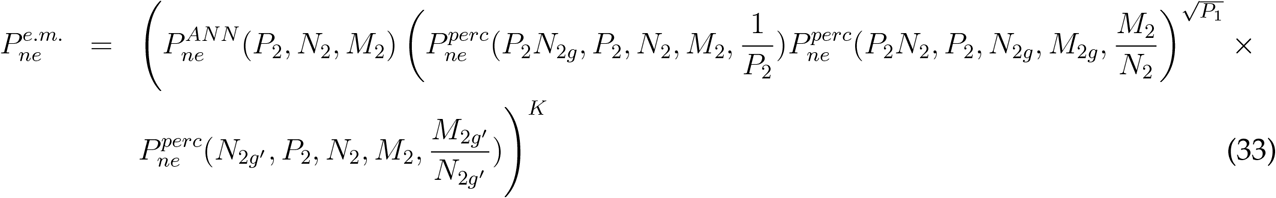

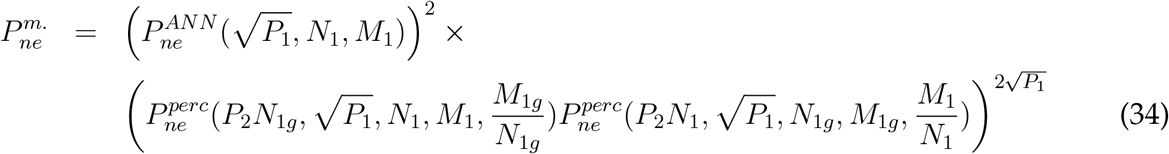

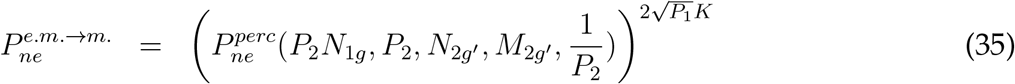

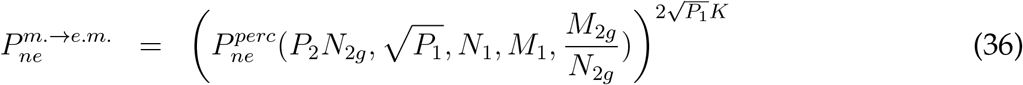

In order to obtain the dotted-dashed curve in Fig. 4e, we considered a special structure for transitions in machine and external memory transitions such that it makes sense to use a compositional representation for them. In our example, half of the cell symbols are associated with transitions of a subset of *P*_1*/*2_ machine states, and the other half of the cell symbols are associated with transitions of the complementary subset of *P*_1*/*2_ machine states. Symbol transitions also follow the same structure, with transitions from the first half of cell symbols depending only on the state of one of the machine NLSC and transitions from the second half of cell symbols depending only on the state of the other machine NLSC. Formally, we have *P*_1_ machine states given by couples *Q*(*t*) = (*Q*^1^(*t*), *Q*^2^(*t*)) with *Q*^1^(*t*) ∈ *𝒯*^1^,*Q*^2^(*t*) ∈ *𝒯* ^2^ with 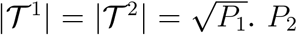 symbols taken from the alphabet 𝒩 𝒯 = 𝒩 𝒯_1_ ∪ 𝒩 𝒯_2_ with |𝒩 𝒯_1_ | = | 𝒩 𝒯_2_ | = *P*_2_*/*2. For a symbol *S*(*t*) ∈ 𝒩 𝒯_*k*_, we have *S*(*t* + 1) = *F*_*k*_(*Q*^*k*^(*t*), *S*(*t*)) and *Q*^*k*^(*t* + 1) = *G*_*k*_(*Q*^*k*^(*t*), *S*(*t*)) and *Q*^*j*≠ *k*^(*t* + 1) = *Q*^*j*≠*k*^(*t*).

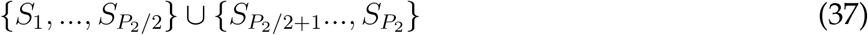

With this constraint on transitions, the number of transition gate neurons in the two machine NLSC and the external memory NLSC can be drastically reduced because we do not need to sub-divide each gates into the 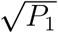 subpopulations. *N*_*min*_ is obtained by minimizing 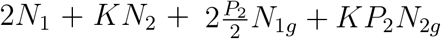 under the constraint 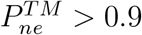 replacing equations **(26), (27)** of **(25)** with

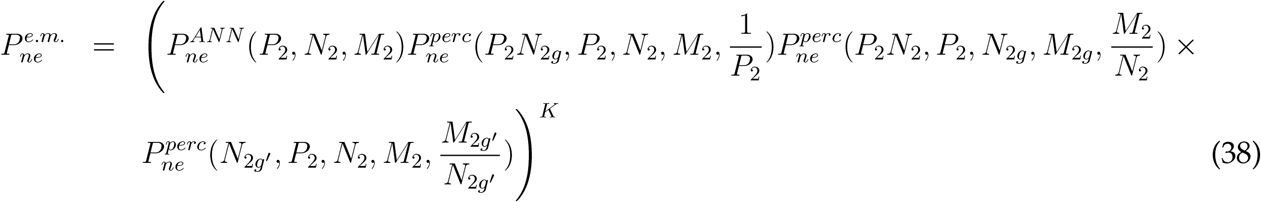

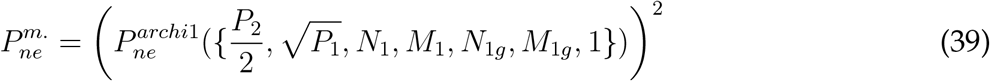

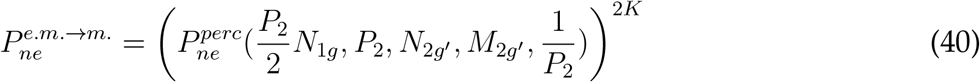

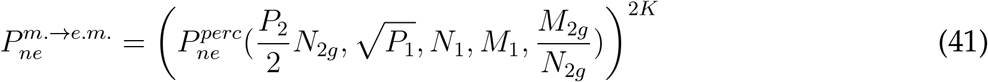

#### 6.1.7 Sequence representations and tree-structures

We consider an environment that produces sequences of length *L* over an alphabet of *D* symbols, with *D* ≫ 1 (10, 000 words) and *L* = *O*(1) (sentence length). We look for a machine that will transform the *D*^*L*^ possible sequential inputs into *D*^*L*^ stable states (meaning of sequences). This can be done straightforwardly by the FSA pictured in SI Fig.6f, which relies on memory states structured into a tree. The NLSC implementing this FSA has 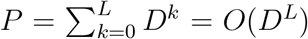 dictionary patterns (SI Fig.6g), and the minimal number of neurons associated to this type of FSA are expected to scale as 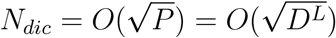 and 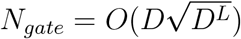 (SI section 6.1.2).

Another possible circuit would buffer the symbols of each sentence into *L* temporary slots implemented by *L* dictionaries with *D* states (SI Fig.6h). This reduces the complexity of the FSA (only *P* = *D*^*L*^ dictionary states), at the price of increasing the complexity of inputs to the FSA. Although the number of neurons required to implement the FSA dictionary is smaller, in the limit *D* ≫ 1 the total number of neurons scales the same way 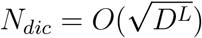 compared to the previous implementation. To ensure proper categorization gate neurons have to mix inputs from the slots and from the FSA dictionary to point towards the correct category. This job is made more difficult by the fact that the *D*^*L*^ possible inputs to the FSA are represented in the *L* slots by non-linearly separable neural patterns. Linearly separable representations for the inputs can be performed by *N*_*gate*_ = *O*(*D*^*L*^). Although we do not prove that this number can not be lowered we still expect it to be larger than 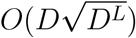, the number of gate neurons used in the neural architecture without memory buffer. This argument could be improved by extending our theory for NLSC to the case *P*_*c*_ = *O*(*P*) (SI section 6.1.2) to compute the minimal number of gate neurons required for this last implementation.

We now consider the case where sequences are structured into chunks: high-level sequences of length *L* are composed on an alphabet of *D* chunks, or non-terminal symbols (SI Section 6.6); each chunk is associated with a sequence of length *l* onto *d* terminal symbols. To categorize these sequences, a naive neural implementation ignoring the chunk structure as the one discussed above will lead to number of neurons scaling as 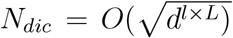 and 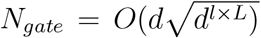. Instead of this naive implementation, we can break down the NLSC implementing the naive FSA into two NLSC (SI Fig.6h) and get rid off the *l × L* product: 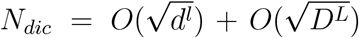 and 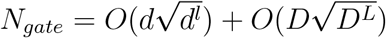.

### 6.2 Low-rank networks

We trained low-rank networks to perform the task of Fig. 1a. To do so we initialized the connectivity matrix as 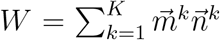 such that recurrently generated activity lies in a subspace of dimension *K* (81) and minimized the loss by updating the parameters 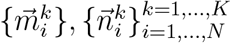, where *N* = *N*_*dic*_ + *N*_*gate*_ is the total number of neurons. Connectivity vectors are structured into populations (78) using the support vectors 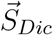 and 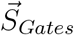 with non-zero entries at 1 on dictionary and gate neurons respectively.

We used a rank *K* = 7 matrix generating the five dictionary states *A, A*^′^, *B, C, C*^′^ and the two states of the external memory *hA, hA*^′^:

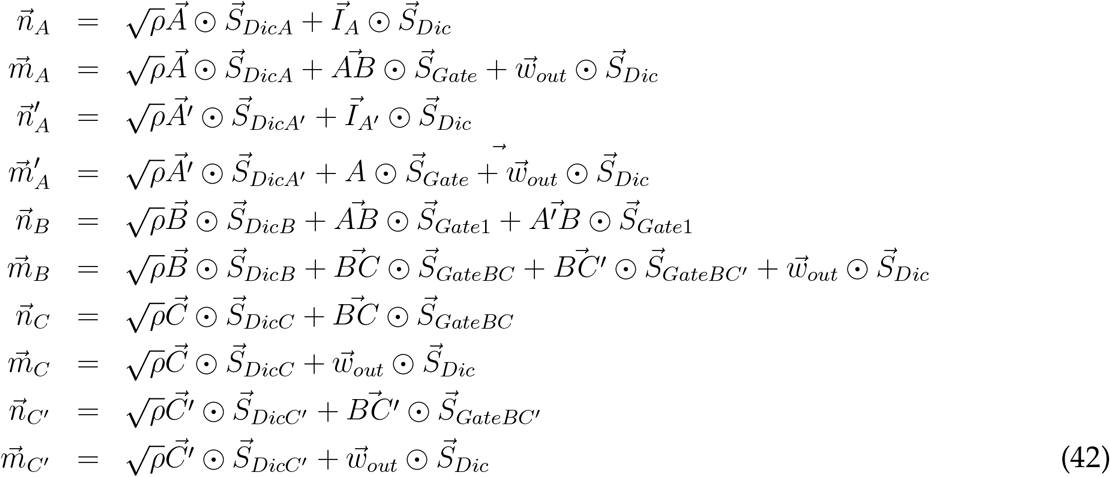

and

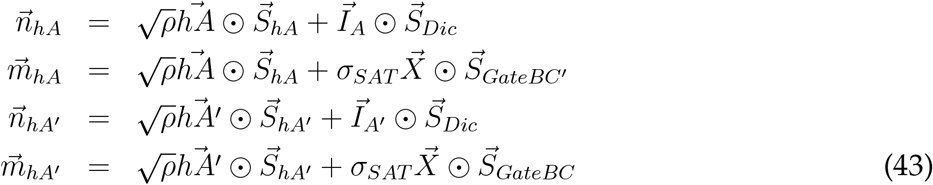

Vectors 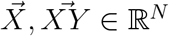 are Gaussian vectors with entries drawn independently from the normalized centered Gaussian distribution. Seven populations, defined with support vectors 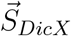 having non-zero entries at 1 on dictionary neurons of population X, support seven independent fixed points by taking *ρ >* 1 such that 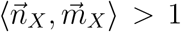. ⨀ is the Hadamard product performing entry-wise multiplication. Vectors 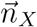 decide which inputs are picked up by recurrent activity. In order for the two cognitive variables *κ*_*A*_ and *κ*_*hA*_ to activate in response to one of the two initializing stimuli, we take 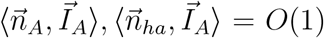, with 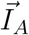 a Gaussian sensory input vectors. Vectors 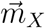 decide in which direction recurrent currents are generated. We take 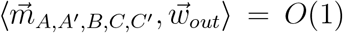 for generating the output, and 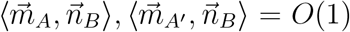 to transfer activations of *κ*_*A*_ or *κ*_*A*_′ into activation of *κ*_*B*_ upon presentation of the go cue, which changes the gain of gate neurons from 0 to 1 through the input 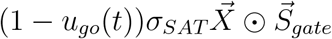 with *σ*_*SAT*_ ≫ 1. We also have saturating currents (*σ*_*SAT*_ ≫ 1) generated by 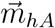 and 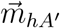 on the two gate populations supported by 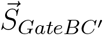 and 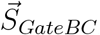, to control transitions from *B* to *C* and from *B* to *C*^′^.

Equations over cognitive variables 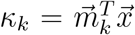 are obtained by projecting the dynamical equation **(1)** onto directions 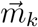 leading to equations **(4)** (78) with

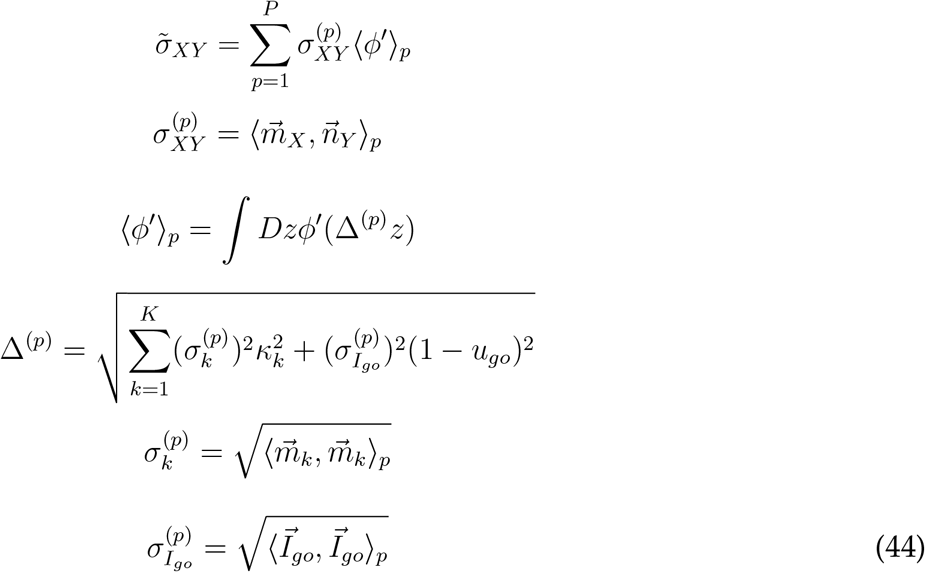

where the scalar products ⟨.,. *⟩*_*p*_ are taken over neurons belonging to population *p*.

### 6.3 Pseudo-code for foraging task

~~~
*# initializations*
nb_paths = 3
P = 10
*# each of the 3 paths is defined as a vector of P subsequent non-
repeating positions*
paths = permutations[nb_paths,P]
*# vector of size P with entries drawn as 0 (no food) or 1 (food) to
initialize reward map*
reward_map = random[0,1,P]
start_position = 1
hunger_level = 3
*# variable holding the actual position of the animal*
current_location = start_position
*# variable holding the virtual position of the animal*
virtual_location = start_position
**while** hunger_level > 0:
       *# planing step:*
       **for** i **in range**(nb_paths):
               *# variable holding the distance between the actual position of
               the animal and the next available food location on path i*.
               count_first_encounter[i] = 10
               reward_found = 0
               **for** j **in range**(P):
                     virtual_location = paths[i](virtual_location)
                     reward_at_virtual_location = reward_map[virtual_location]
                     **if** (reward_at_virtual_location == 1) **and** (reward_found ==
                          0):
                             count_first_encounter[i] = j
                             reward_found = 1
       *# decision step*
       chosen_path = argmin(count_first_encounter[:])
       nb_steps = **min**(count_first_encounter[:])
       *# traveling step*
       **for** j **in range**(nb_steps):
             current_location = paths[chosen_path](current_location)
       *# food consumption step*
       reward_map[current_location] = 0
       virtual_location = current_location
       hunger_level = hunger_level - 1
~~~

### 6.4 Assembling NLSC for the foraging task

The head circuit is a NLSC with a 10-states dictionary each encoding one of the 10 positions. This dictionary is associated with three gate populations, each encoding a permutation, or a path, across the 10 locations. Wiring of the head circuit reflects information encoding in the parameters contained in **paths** initialized at the beginning of the pseudo-code. Updates of the variables **current location** and **vritual location** are implemented by dictionary state transitions in the head NLSC.

The external memory implements the **reward map** variable. It is composed of 10 NLSC, one per location (Fig. 5d, green square). Each is composed of a 2-states dictionary encoding the absence or presence of reward at the corresponding location, together with a single gate population activated to update reward value, e.g. when food is consumed at this location. As explained in Method 5.2.4 the connectivity from the dictionary’s head circuit to the gates of the external memory are first initialized: a cell assembly encoding location *X* in the dictionary inhibits all 10 gate populations of the external memory, but the one associated to the NLSC encoding value at position X. This connectivity is then fine-tuned by training the assembled network to update, upon activation of an external go cue, the value of the external memory NLSC-X only when the head circuit is in state X. In addition to these *transition* gates bi-directionaly coupled with the dictionary and implementing transitions between dictionary states, these NLSC have *output* gates that only receive from the dictionary and that are modulated by the head circuit to decide from which of these 10 NLSC the machine is focusing its attention.

What we considered here as the machine operating on the external memory through the head circuit is composed of multiple NLSC with different functions (Fig. 5d, blue square). We first present NLSC that implement specific operations, and then NLSC that coordinate these operations. To each path is associated a counter memorizing the distance from food on this path (**count first encounter** variable). Each counter is a NLSC with a 10-states dictionary encoding distances from 1 to 10 and a single gate population implementing a single permutation allowing to increment the counter. As for the external memory, these NLSC have *output* gates that provide, after the planing steps, the number of locations to go through to reach reward.

These transitions in the counters only occur when a binary memory slot signals that no reward has yet been encountered during planing a path. This memory, carried by the **reward found** variable in the pseudo-code, is implemented by a NLSC with a 2-states dictionary, encoding whether or not a reward has been encountered (Fig. 5g). A single gate population activates to trigger transitions between these two states, with e.g. an update when a location with reward is encountered (Fig. 5g).

When the three paths have been planed, a decision circuit reads from the 3 counters to select which path will actually be taken. This decision circuit is a NLSC with a dictionary state composed of a neutral state and 3 states each encoding a potential chosen path. A gate neuron population receiving inputs from the 3 NLSC counters has its connectivity with the dictionary trained to perform the *argmax* operation, transiting from the neutral state to the state associated to the path *P* ^∗^ with the smallest counter value, updating the variable **chosen path**. Once such a state is reached, it releases inhibition on the gate population corresponding to the chosen path in the head NLSC such that the agent travels on path *P* ^∗^ through *K*_*P*_ ∗ locations.

Reaching reward, food consumption is modeled by switching to an eat-state in the 2-states dictionary of a NLSC. Upon this switch, inhibition is released on the gate population of another NLSC. This hunger level NLSC has a 5-states dictionary each associated with a possible hunger-level of the agent (**hunger level** variable). The dictionary state associated with hunger level 0 inhibits all gate populations in the neural architecture, terminating the **while** loop in the pseudo-code. The gate population triggers transition towards a state encoding the next lower hunger levels, corresponding to the line of the pseudo-code where the **hunger level** variable is updated.

The **inner for-loop** in the planing step of the program (SI Section 6.3) is implemented by a NLSC with a 10-states dictionary. This dictionary is associated with a gate population implementing a single permutation of these 10 states. Before planing each path, the dictionary of this NLSC is in state 1, the gate population activates each time the head circuit is updated such that after going through a path, it ends up in state 10. The cell assembly associated to this final state informs, through its connectivity to a gate population, a coordinating NLSC to plan on another path or to move on to the decision step. This coordinating NLSC is composed of a 5-states dictionary. 3 states are associated with planing the 3 paths (**outer for-loop**), one state is associated with the decision and traveling steps (when the corresponding cell assembly is activated it allows for transitions in the decision making NLSC and enables movements of the agent) and one state is associated with the food consumption step (when the corresponding cell assembly is activated it allows for transitions in the eat, reward map and hunger-level NLSC).

### 6.5 Automata

In order to support the claims made throughout the main text, we recap on what is presented in (25) about automata and computation.

#### 6.5.1 Finite-state automata

A finite-state automaton (FSA) or finite-state machine is composed of a finite number of states *Q* ∈ {*Q*_1_, …., *Q*_*P*_}, at each time step it can receive a stimulus *S* ∈ {*S*_1_, …, *S*_*K*_}, or symbol, from its environment. In response to that stimulus it updates its state *Q*(*t*) to *Q*(*t* + 1) and might produce a response *R*(*t* + 1):

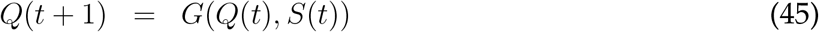

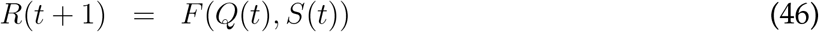

Below we give specific examples of FSA to illustrate the work exposed here.

FSA with *P* states can be used to build a head-direction system with each state *Q*_*k*_ representing an angular position 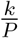 360^□^ (24). At any time, the environment of this system (e.g. the vestibular part of the nervous system of the animal) provides one of two stimuli *S* ∈ {−1, +1}, indicating if the animal’s body has rotated counter-clock wise or rotated clock wise. Such automaton can be referred to as a categorizer in the sense that it categorizes all the possible rotation sequences (e.g. +1, −1 or +1, +1, +1, −1, +1, −1) into one of *P* classes each representing a current head-orientation with respect to some arbitrary landmark.

A peculiar class of FSA is the class of autonomous FSA, or sequential machines ((24), Fig.4a). They take only a single symbol as input, instructing the automaton to move on to its next state. A common example is the one of a traffic light that switches states each time a timer signal is ON, with each state associated to the production of a color.

Another class of automata is the class of acceptor or recognizer. Upon receiving a string of symbols taken from an alphabet, it tells, with a final state, whether the string has been generated by a given regular language L (accept final state), or not (reject final state) (SI section 6.6).

Note that we have used two different graphical representations for FSA. In (24) and in SI Fig. 1h,l, we represented each state by an hexagon together with labeled arrows telling which input would trigger which transition between states. In the main text here we used a more compact representation in which the entire FSA is represented by a box, the front-side of which displays the current state it is in.

#### 6.5.2 Turing machines

Modeling the operations of machines with FSA can be considered incomplete in the sense that reactions of the environment *S*(*t* + 1) in response to the actions of the machine *R*(*t*) are left unspecified. In order to model and define the notion of computation, Turing proposed a minimal model in which FSA are augmented with a head that can read, write and travel on an external memory modeling the environment (Fig. 4c). The tape is split into *K* cells each displaying at time *t* a symbol *S*^*k*^(*t*) ∈ {*S*_1_, …, *S*_*K*_}. If at time *t* the head reads cell *k*_0_(*t*) of the external memory, it leads the machine to i) update its state according to

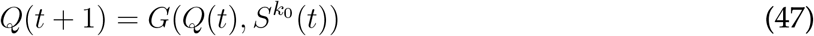

to ii) act upon the external memory in a predictable manner by updating the current cell’s state’s according to a well defined function *F* (.,.)

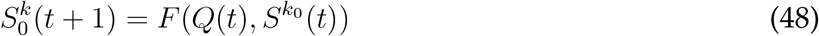

where F and G are independent on *k*_0_, as well as to iii) move the head to another location depending on what has been read

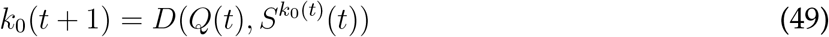

where 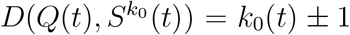 such that the head moves immediately either left or right from the current cell being read. Turing machines are automata composed of such FSA, head and tape.

In our neural instantiation of Turing machines, machine states and tape symbols are carried by cell assemblies in dictionaries while transition tables represented by the functions *G*(.,.), *F* (.,.) and *D*(.,.) are instantiated by linearly separable representations in gate neurons and specific wiring between dictionary and gate neurons.

#### 6.5.3 Other automata

While FSA alone are limited in the procedures they can implement, it is commonly postulated (Church-Turing thesis) that any procedure that can be well-defined can be implemented by a Turing machine, provided that the tape can grow an infinite number of cells. This is done by specifying the alphabets of the machine (*Q*^′^*s*) and tape-cells (S^′^*s*), as well as the transition functions *G*(.,.), *F* (.,.) and *D*(.,.). Turing machines can be extended with e.g. multiple tapes or in other way, but the obtained automata do not gain in computational power. Otherwise, other automata can be defined as restrictions from Turing machines. Different classes of automata maps onto levels of the Chomsky’s hierarchy as explained below.

For instance, linear-bounded automata are Turing machines with a non-infinite tape. This class of automata has been shown to contain acceptors for languages generated by context-sensitive grammars. Pushed-down automata can be seen as Turing machine with a single-cell tape with restrictions on the way symbols are updated on this cell. This class of automata has been shown to contain acceptors for languages produced from context-free grammars. Finite-state automata are Turing machines without tape (or with a tape read from left to right without back and forth), and this class contains acceptors for languages produced from regular grammars.

### 6.6 Formal languages, generative grammar and automata

**Formal languages**, such as those defined by generative grammar, are sets of words obtained by forming strings of symbols taken from an alphabet 𝒯 of terminal symbol size *P*. Grammatical or production rules use another alphabet of *non-terminal* symbols 𝒩𝒯. Applying these rules in succession leads to the production of strings of terminal symbols that belong to a specific language. In their most general forms they take the form *α* → *β*, with *α* and *β* that can be any string formed by terminal and non-terminal symbols. Languages generated by such a rule are called recursively enumerable and constitute the highest-level class of languages of the Chomsky’s hierarchy.

Other lower-level classes of languages are defined based on restrictions on this form of rule. For instance, regular languages are produced by rules in which only a single non-terminal symbol is allowed on both the left and right hand-side of the arrows. An example is illustrated by the language {*a*^*k*^*bc*^*k*^, *k* ∈ ℕ} composed of all strings composed of *k* successive *a, b* and *k* successive *c*, on an alphabet of terminal symbol 𝒯 = {*a, b, c*}. It can be produced by applying the following rules defined with the non-terminal symbols 𝒩𝒯 = {*S, Q*_1_, *Q*_2_, *Q*_3_, *Stop*}:

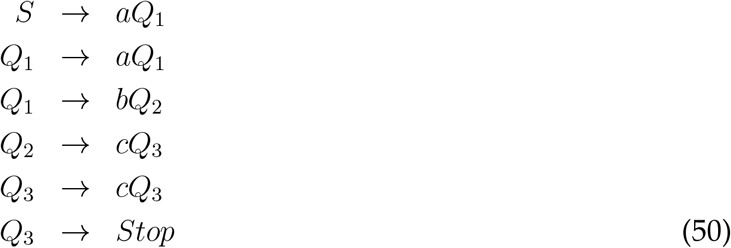

#### Relationship between Chomsky’s hierarchy and automata

Languages belonging to a specific class are those that can be recognized by automata with specific computational abilities.

Regular languages are defined by production rules of the form *A* → *aB*, with *A* or *B* standing for one of the non-terminal symbol and *a* for one of the terminal symbol. They are recognized by finite-state automata. For instance the language {*a*^*k*^*bc*^*k*^, *k* ∈ ℕ} is recognized by the FSA depicted in Fig.

Context-free languages are defined by production rules of the form *A* → *a*, with *A* a single non-terminal symbol and *a* here stands for any string of both terminal and non-terminal symbols. They are recognized by push-down automata (SI Fig. 1c). A push-down automata is a FSA augmented with an external memory constituted by a stack of symbol. The identity of the first symbol on top of the stack determines the mapping between state-transitions and inputs in the FSA. At each time step, the first symbol is either popped from the stack, leading the second symbol to be on top, or a new symbol is put on top of the stack, leading the first symbol to become the second one.

Context-sensitive languages are defined by production rules of the form *αAβ* → *αγβ*, with *A* a single non-terminal symbol and *α, β, γ* stands for any string of both terminal and non-terminal symbols. They are recognized by linear-bounded automata: Turing machines with a tape whose number of memory cells is finite and proportional to the number of cells on which the input to be computed is written.

Unrestricted languages are defined by production rules of the form *α* → *β*, with *α* and *β* that can be any string formed by terminal and non-terminal symbols. They are recognized by Turing machine with tapes with an infinite number of memory cells.

Below we give examples of production rules that describe some of the behavioral tasks that we consider in this work. An example is the following set of rules {*R*_*k*_} over 𝒯 = {*a, b*, …, *z*} and 𝒩𝒯 = {*Start, Stop, A, B*, …, *Z*}

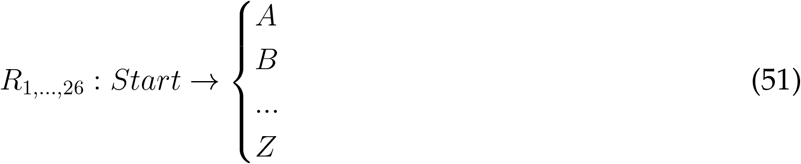

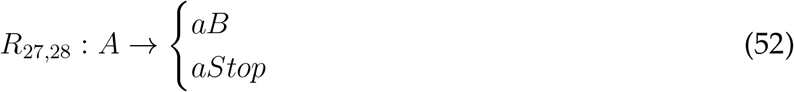

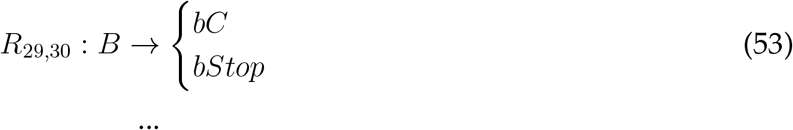

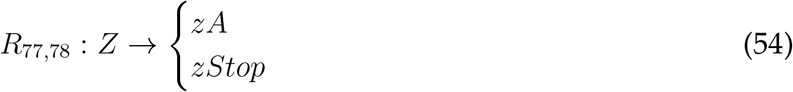

These alphabets and production rules in finite number define the language L as an infinite subset of the set of all the possible strings that could be composed from the alphabet 𝒯, namely the subset of strings that are arranged in alphabetical order ((24), Fig.2b). Every string in L can be printed by application of the rules, e.g. string *abc* is obtained by application of *R*_1_, *R*_27_, *R*_29_ and *R*_32_.

The above language is a regular language, with only a single non-terminal symbol allowed on the right-hand side of the arrows. Allowing multiple non-terminal symbols on the right hand-side, the strings produced from applications of such grammatical rules can exhibit long-distance dependencies. For instance in the following example generating the finite set of sequences of Fig. 1a with an alphabet of terminal symbols 𝒯 = {*a, a*^′^, *b, c, c*^′^} and non-terminal symbols 𝒩𝒯 = {*Start, Stop, S*1, *S*2, *B, C*}

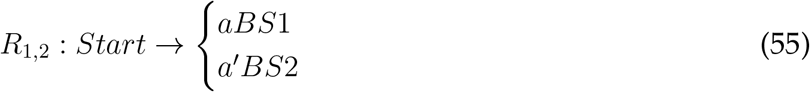

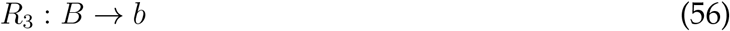

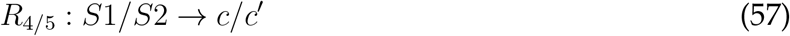

for a finite set of sequences it is always possible to propose rules with regular-language restrictions (e.g. *S* → *abc/*a^′^*bc*^′^). By contrast, in order to generate the infinite set of sequences with long-distance dependencies that we considered in SI Fig. 3a, a compact description of all the possible sequences is given by context-sensitive production rules. Context-sensitive production rules can have multiple symbols on the left-hand side such as

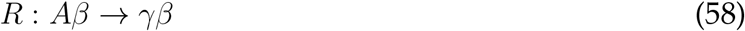

where *β* and *γ* are strings of terminal and non-terminal symbols, with *γ* non-empty. The symbol A is thus replaced depending on its surrounding context *β*. For our example, a language with all alphabetical ordered sequences starting from odd letters (A, C, …) and counter-alphabetical ordered sequences starting from even letters (B,D,…), the production rules are the following

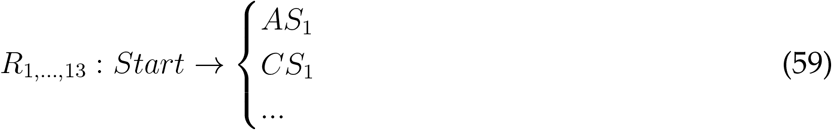

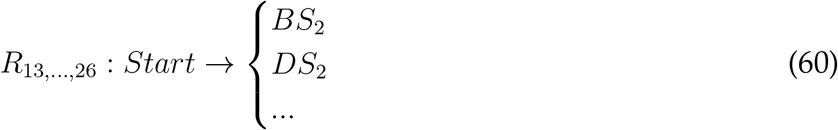

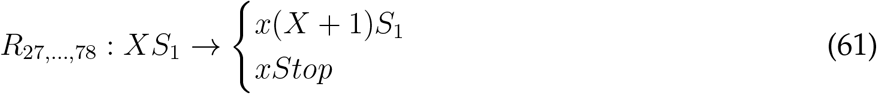

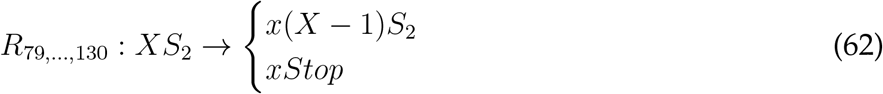

with alphabets 𝒯 = {*a, b*, …, *z*} and 𝒩𝒯 = {*Start, Stop, A, B*, …, *Z, S*_1_, *S*_2_}. These rules can easily be generalized to generate sequences organized hierarchically (Fig. 2a), by introducing non-terminal symbols for chunks at multiple levels. By contrast, writing generative grammar rules that produce sequences with an algebraic pattern structure (23) such as XYX (Fig. 3a) is cumbersome, owing to the requirement that the two X’s should lead to printing the same symbol. Possible context-sensitive rules are the following:

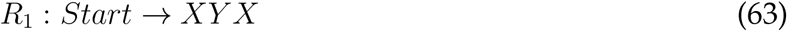

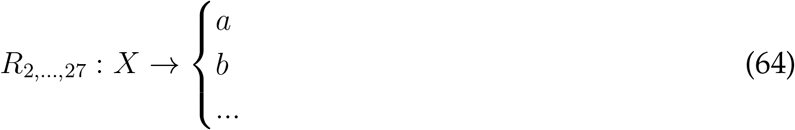

when the context on the left of X is empty, while if this context is non-empty X yields

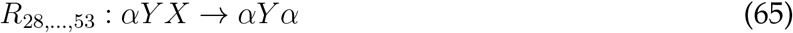

for *α* ∈ 𝒯. And,

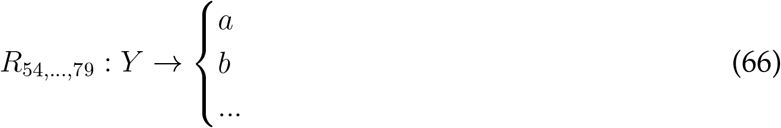

We needed to introduce many particular rules for the expression of the last X (*R*_28,…,53_), making the generative grammar formalism not well suited to compactly capture this type of temporal structures.

### 6.7 Relationship with natural language and tree structures

We find that the primitive temporal structures outlined in (23) maps more naturally onto properties of neural architectures compared to the levels of complexity defined by Chomsky’s hierarchy. For instance when going from regular to context-free languages, the associated automata are FSA and push-down automata. Pushed-down automata are composed of a FSA reading on a single-cell external memory, as is the case in the neural architecture proposed for sequences organized into chunks. However the functioning of this external memory, which works as a stack of symbols where the top symbol can be popped or replaced by a new one, is more difficult to map onto neural network operations compared to the single cell-external memory composed of a NLSC.

Beyond sequences organized into chunks or algebraic patterns, cognitive science studies outlined that high-level cognitive processes such as natural language relies on tree structures (23). Following physical constraints on neural networks, we found it more natural to jump from neural architectures for sequences organized into chunks or algebraic patterns, to neural Turing machines associated with the running of sensory-motor programs. In our framework, tree-like mental constructions appear when trying to build a circuit that receives strings of words and maps those into stable meaning representations (SI Section 6.1.7). Building such meaning representations requires to use intermediary neural representations that evolve on a tree structure (SI Fig.6f). Tree-structures can be repeated at multiple levels of abstractions if sequences to be mapped are organized into chunks (SI Fig.6h). Within this setup, we compared a scenario where each new incoming symbol is merged with the current neural representation (SI Fig.6g right), with a different scenario where incoming symbols are buffered before being merged (SI Fig.6g). We found the first scenario to be more economical in terms of number of neurons (SI Section 6.1.7). We wonder whether this physical constraint on neural architectures could be related to the MERGE operation that has been postulated to be the fundamental mental operation underlying natural language (23).

### 6.8 Supplementary discussion

#### 6.8.1 Relationship with previous theoretical works

##### Statistical physics of neural networks

The elementary circuit is composed of a dictionary in which cell assemblies can be activated thanks to positive feedback loops in its connectivity. The dictionary is augmented with gates allowing to trigger transitions between cell assemblies. We have used tools from statistical physics to characterize the NLSC. From a dynamic-mean-field point of view (82; 78), cell assemblies are associated with cognitive variables, the gain of populations of gate neurons regulate functional interactions between cognitive variables (SI Fig.4) (78). We have leveraged storage capacity calculation techniques originally applied to perceptrons (29; 30), Hopfield-like models (83; 48; 82; 31) and attractor neural networks in general (84; 85; 86), to provide a quantitative characterization of NLSC with minimal number of neurons. This minimal number of neurons is obtained when minimizing interferences between auto-associative loops of the dictionary and the hetero-associative connectivities with gate neurons (Fig.1h,i, SI Section 6.1). This is done by tuning synaptic connectivity probabilities, which are controlled by the coding levels of neural patterns under the Hebbian learning scenario considered here. As for previous works (48; 80), we found that the maximal number of stable states and permutations of states are reached for small coding levels, for which Hebbian-type plasticity leads to an optimal use of neural ressources (87).

##### Sequences in neural networks

Focusing on temporal structures, our work departs from a previous work interested in simple transitions between attractor states (88) or from storage capacity studies of networks implementing unstructured sequences either with direct hetero-associative connectivities (89; 90; 91; 92; 93) or gated-hetero-associative connectivity as used here (94). Previous works proposed networks producing or recognizing sequences with hierarchical temporal structures (95; 96). Gate neurons similar to the one we introduced have also been proposed to model sub-cortical circuits and to confer flexibility in generating sequential behaviors (21; 54).

##### Neural networks population structure and temporal structure

Here we set out to systematically study temporal structures, and described how NLSC can be combined to produce sequences with various forms of temporal structures. Previous works studied context-dependent computations (34) and found that external contextual inputs could reconfigure functional interactions between cognitive variables and inputs, provided that networks are structured into populations (78). Some population implementing attractor dynamics similar to dictionaries and other populations, gain-modulated by the contextual inputs, similar to gate neurons. Here we treated input from dictionary states onto gate neurons as contextual inputs, such that, from the dynamic-mean-field point of view, cognitive variables in one dictionary controls the functional interactions between cognitive variables of another dictionary (Method 5.4), with bi-directional communications between two levels allowing structured sequences to unfold (SI Fig.4). Storage capacity calculations coupled with a minimization principle have allowed us to relate the population structure of the architectures with the behavior they implement. We thus provide a theoretical background that supports our previous observations regarding the emergence of population structure in ANN (78): gate neurons of each NLSC segregate into populations to mix current dictionary states and external inputs into linearly separable representations ((24), Fig.5), while neural architectures are segregated into NLSC depending on computational demand (Fig.4e).

##### Computations through neural dynamics perspective

The computations performed by Hopfield or attractor networks are obtained by convergence towards fixed points. Dynamical features such as fixed points, stable and unstable, or limit cycles have been proposed to underlie other forms of autonomous neural computations. This point of view is often referred to as computation through neural dynamics (46). Computations on time varying external inputs have also been described within this framework, both for inputs aligned with the network recurrent structure, allowing temporal integration (34) or recognition of sequences of inputs (97), as well as for inputs orthogonal to the network recurrent structure, allowing contextual inputs to reconfigure network’s dynamics (34; 46; 78). We have shown in (78) how such external inputs, by modulating the gains of specific populations of neurons, allow to reconfigure the interactions between aligned inputs and cognitive variables sustained by recurrent activity. We have shown how external go cues can also reconfigure interactions among cognitive variables supported by the recurrent connectivity of NLSC’s dictionaries, by controlling the gains of specific populations of gate neurons ((24) SI Fig.1, Method 5.4). We have also shown how the activation of high-level cognitive variables can play the same role as these external inputs and reconfigure interactions between lower-level cognitive variables (SI Fig.4). These modulatory interactions among cognitive variables are key to organizing communication among NLSC so as to endow neural architectures with elaborated computational abilities.

#### 6.8.2 Neural networks as automata

##### Functioning of neural Turing machines

In order to provide a well defined notion of temporal structure, we interpreted cognitive variables and their associated cell assembly activations as automata states. This has allowed us to interpret various forms of temporal structures as restrictions on the computational abilities of automata. By building Turing machines, we have shown that combining NLSC allows in principle to implement any program, procedure or algorithm. A program is implemented by wiring connectivity between dictionaries and gates in order to have the correct context-dependent transitions in the dictionaries of the external memory, the machine and the head circuits. As input of the program, the external memory is loaded by setting the dictionary of each NLSC into persistent states representing cells’ symbols. Transitions between machine, head and cells states correspond to computation unfolding in time. The output of the program is the final state of the tape or of the machine.

##### Finite neural networks

In practice, the neural Turing machines we introduced have a finite tape as any practical machine. In order to state that any program can be implemented by combining NLSC, we follow classic arguments (25) and assume that new cells can be added to the external memory if the computation requires it. The quantitative theory we developed allows to compute how many neurons are required to add a new cell to a given architecture. We note that adding a new cell requires adding a new NLSC with the correct wiring with the FSA, so as to implement the set of transitions between cell symbols and machine states associated with the program to be implemented. We also note that human working memory is rather limited, more or less seven items, such that looking to build machines with infinite memory might not be particularly relevant to model the functioning of brain circuits.

#### 6.8.3 Relationship with brain circuits

##### Recurrent circuitry and cell assemblies

In dictionaries of NLSC, neural dynamics exhibits attractor states which have been shown to be well suited to encode cognitive variables involved in specific brain computations (13). Early physiological characterizations of brain circuits align well with the notion of attractor neural networks, with persistent activity recorded in various cortical areas during memorization phases of delay-match-to-sample tasks (98; 99; 15; 100). Modern experimental techniques, allowing to perturb neural states in behaving animals, have allowed to directly probe the attracting nature of neural states linked to behavior, for instance in motor-cortex (101) or in fly’s ellipsoid body responsible for the encoding of head-orientation (102). On the anatomy side, the recurrent circuitry of the brain appears well suited to implement attractor states, for instance it has be shown that statistical features of local cortical connectivity matches those of attractor neural network’s connectivity optimizing memory storage (33; 103).

##### Oscillatory activity

There are experimental evidence that local microcircuitry could support autonomous state transitions through oscillations, for instance it has been observed that populations of hippocampal PV cells oscillating at *γ*-frequency update their state at each oscillation cycle (72). Hippocampal replays at *θ*-frequency could also be interpreted as updating the state of a NLSC. It would also be nice to understand how oscillations can coordinate various NLSC involved in running a hierarchy of sensory-motor programs. Along such a line it has been observed that in an attentional system such as the frontal-eye-field, LFP recordings exhibit *β*-oscillations while animals switch their attention at *β*-frequency in a spatial visual attention task (104).

## Supplementary figures

**SI Fig. 1:**
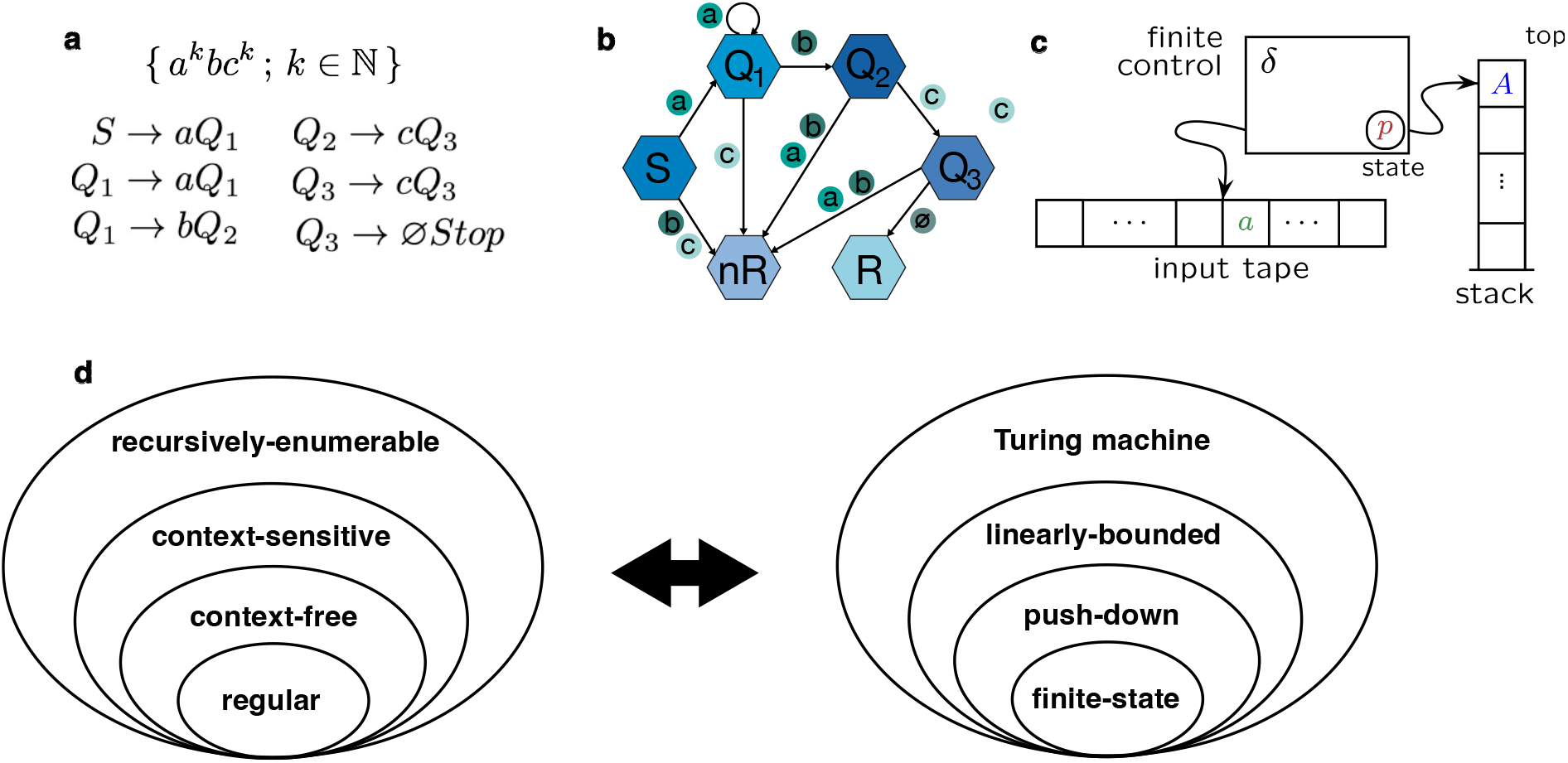
Chomsky’s hierarchy and its equivalence to an automata hierarchy. **a**, Generative grammar rules producing the language {a^k^bc^k^, k ∈ ℕ}. **b**, FSA recognizing this language, the automaton ends up in R, or nR, when the presented string belongs, or not, to the language. **c**, Schematic of a push-down automaton (taken from Wikipedia). They can be used to build recognizer or acceptor for context-free language. The input tape contains a string of terminal symbols and is read from left to right. **d**, The Chomsky’s hierarchy consists of classes of generative grammar rules, each being defined from the previous class by adding a restriction to the form of the rules. At each class of rules corresponds a class of automata. For a given language produced by rules of a given class, one can build a recognizer with an automaton belonging to the corresponding class.

**SI Fig. 2:**
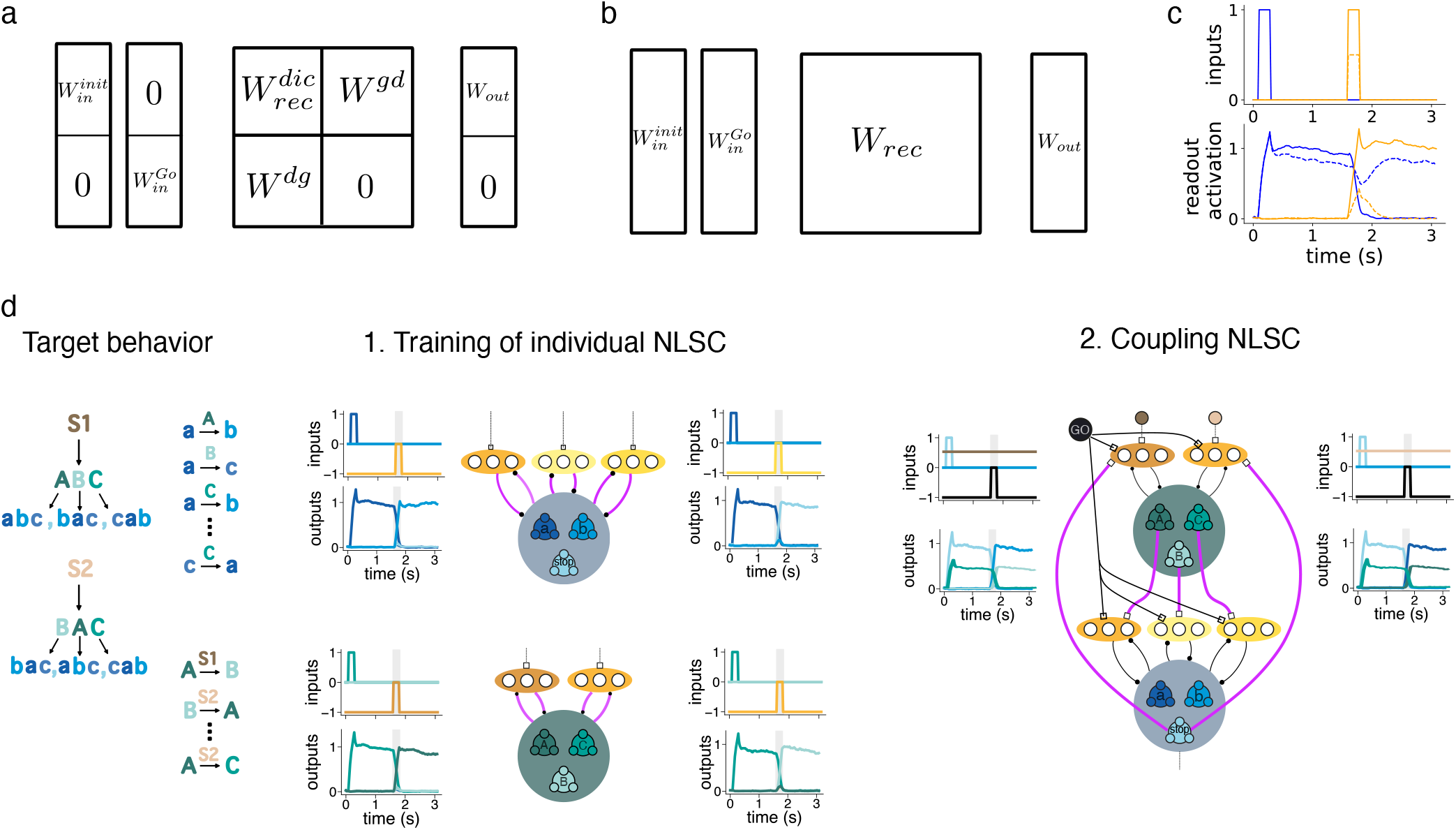
Training schemes for NLSC or for unconstrained ANN. **a**, Schematic of constraints on the connectivity for training NLSC, or **b** unconstrained networks. **c**, Top: activation of two input neurons for two trials (plain and dashed lines) for the dictionary task. Bottom: activations of the two corresponding blue and orange readout neurons for a trained networks in response to these inputs. **d**, Assembling NLSC, illustration on the task of producing sequences organized into chunks. Left: the behavior of interest is split into individual transitions. Middle: individual NLSC are trained to implement relevant transitions. One population of gate neurons is used for each context associated with a permutation of dictionary states. Right: NLSC are coupled to each others by training multiple NLSC to switch their states simultaneously in response to a go cue input (black neuron).

**SI Fig. 3:**
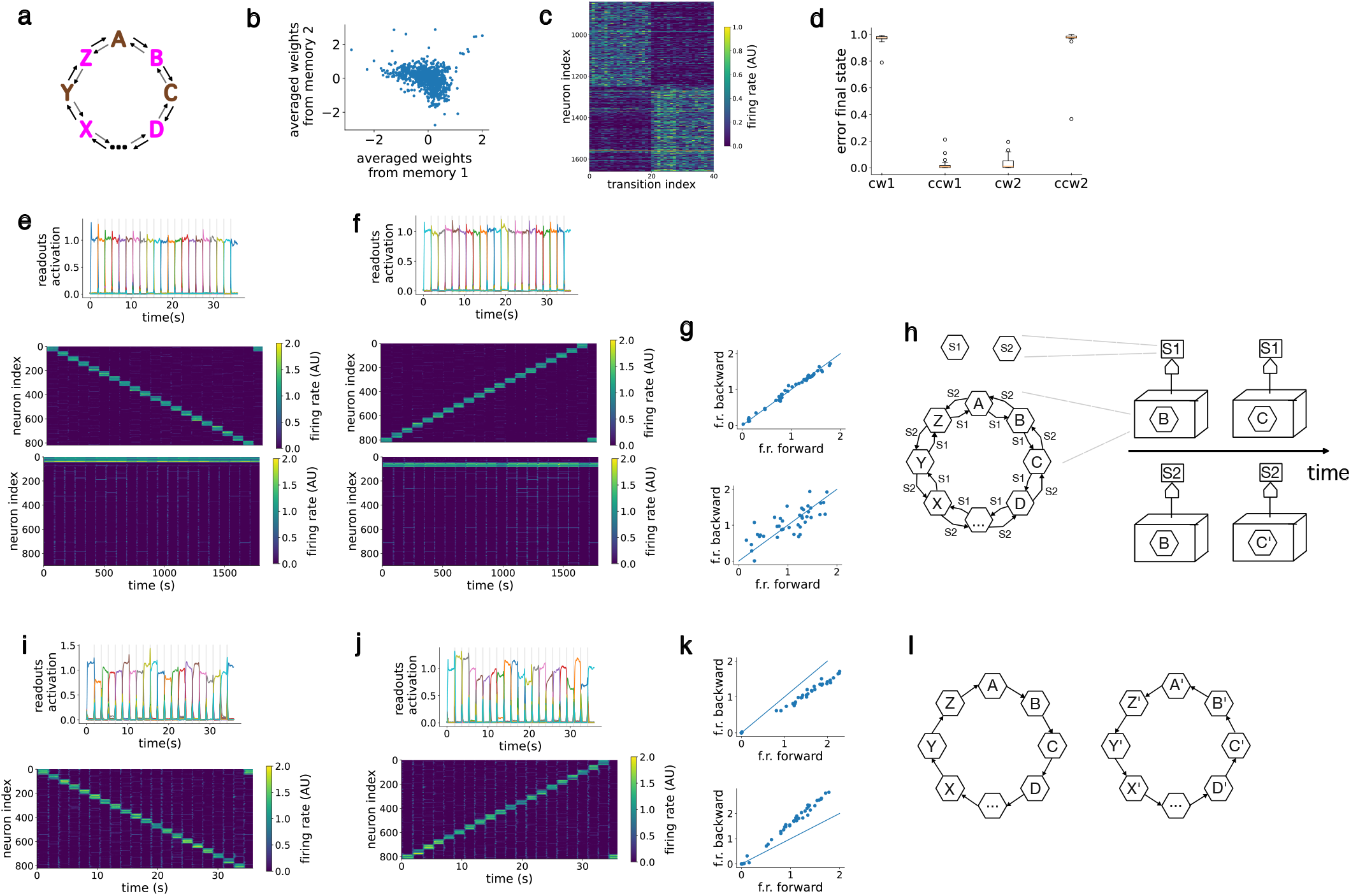
Neural networks for long-distance-dependencies. **a**, Illustration for the tasks with 20 states. **b**, Averaged weights from the two cell assemblies of the external memory onto gate neurons of the NLSC, for the network discussed in Method 5.3.2. **c**, Raster of gate activation patterns for the 2 ∗ 20 transitions. **d**, Results of inactivating the two gate sub-populations in each of the two contexts. **e**, Activity of output (top), dictionary of the NLSC (middle) and external memory and gate neurons (bottom) in the first, **f** second context. **g**, Activation of a cell assembly of dictionary neurons while the network produces a symbol in both contexts. Top and bottom show two examples. **h**, Automata cartoon summarizing the functioning of the NLSC augmented with an external memory. **i-l**, Same as **e**-**h** for an unconstrained networks trained on this task.

**SI Fig. 4:**
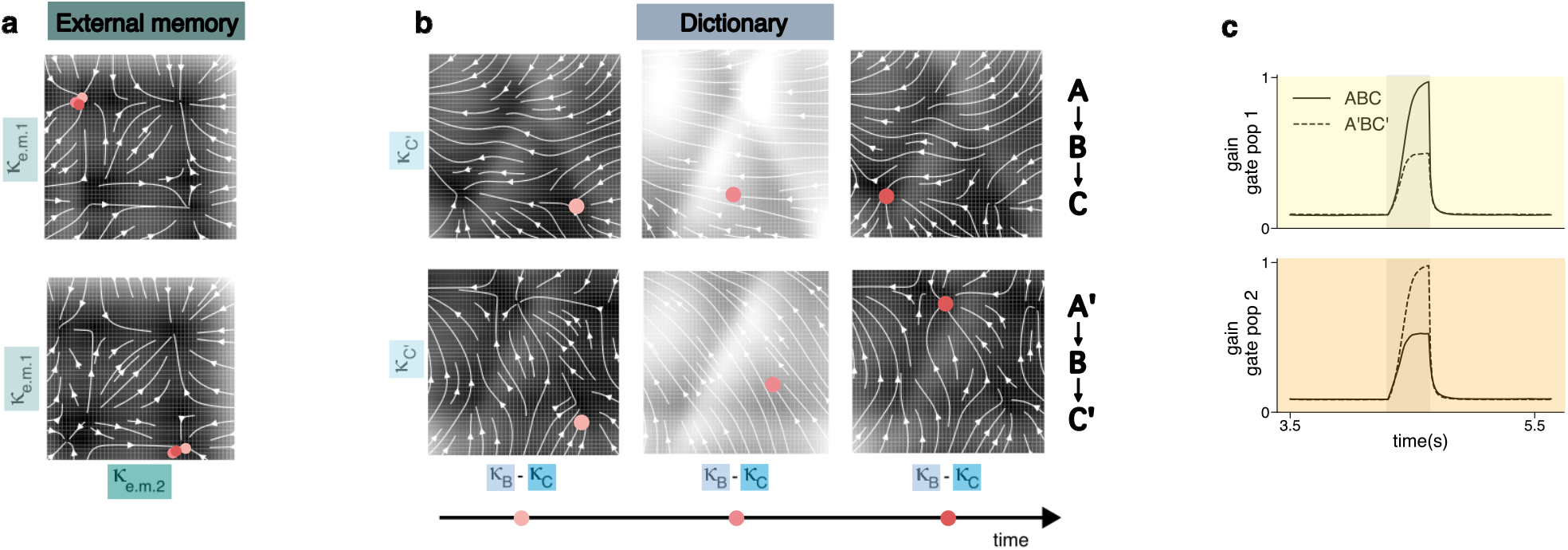
Reduced dynamical systems for NLSC with an external memory solving a task with long-distance dependencies. **a**, Flow-fields depicting the dynamics in the subspace representing the neural activity in the external memory, points of different colors represent neural activity location at different time points throughout sequence production. **b**, Flow-fields in a subspace representing activity in the dictionary of the NLSC, we chose a cut allowing to visualize the three fixed points associated with the production of B, C and C’. **c**, Gain of the two populations of gate neurons routing activity from B to C or B to C’.

**SI Fig. 5:**
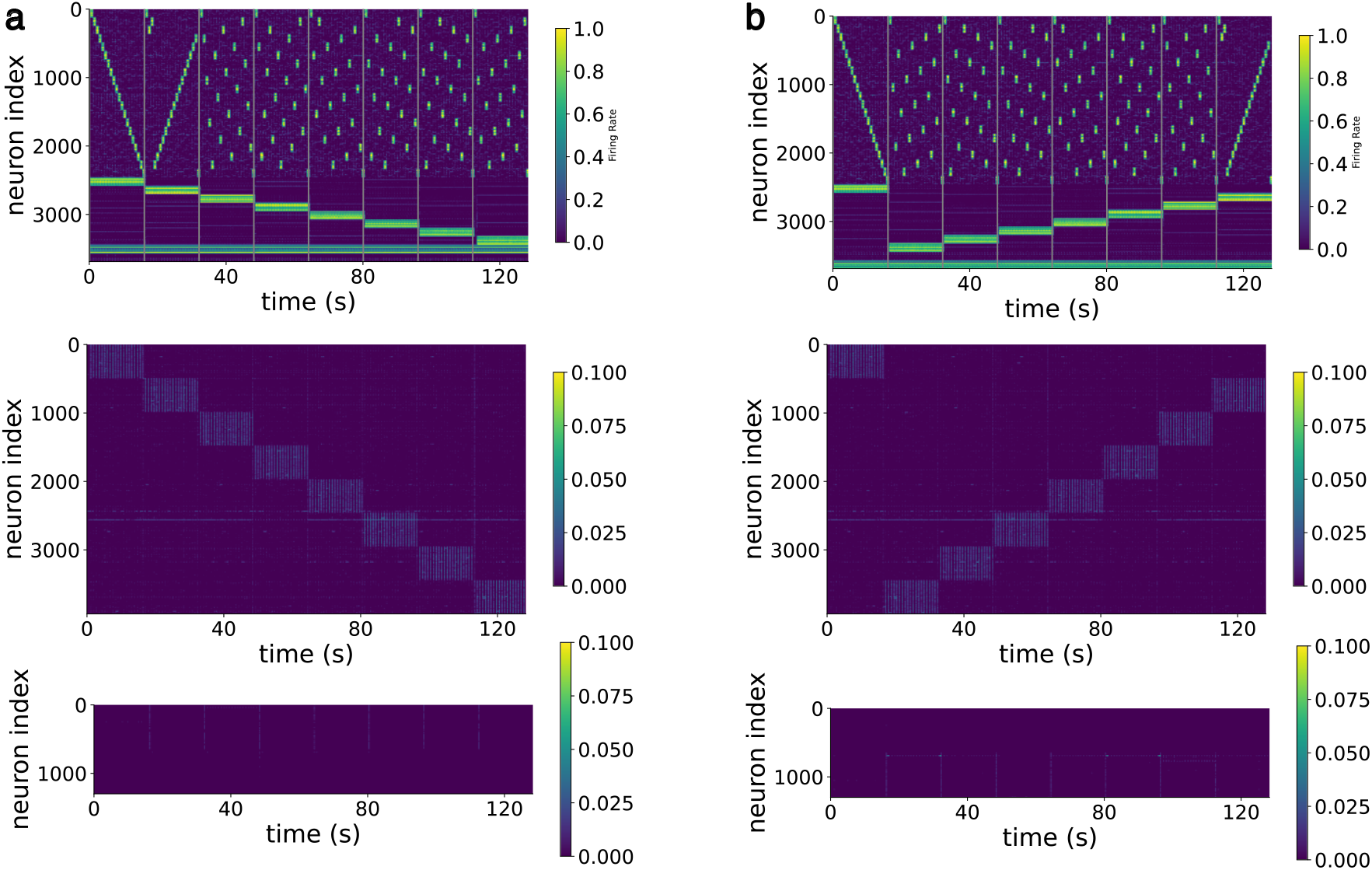
Coupling NLSC. For the architecture presented in Fig. 2b, rasters of activations of dictionary neurons in the three NLSC (top), gate neurons associated with the lowest-level NLSC (middle), gate neurons associated with the high-level NLSC while playing sentence S1 **a** or S2 **b**.

**SI Fig. 6:**
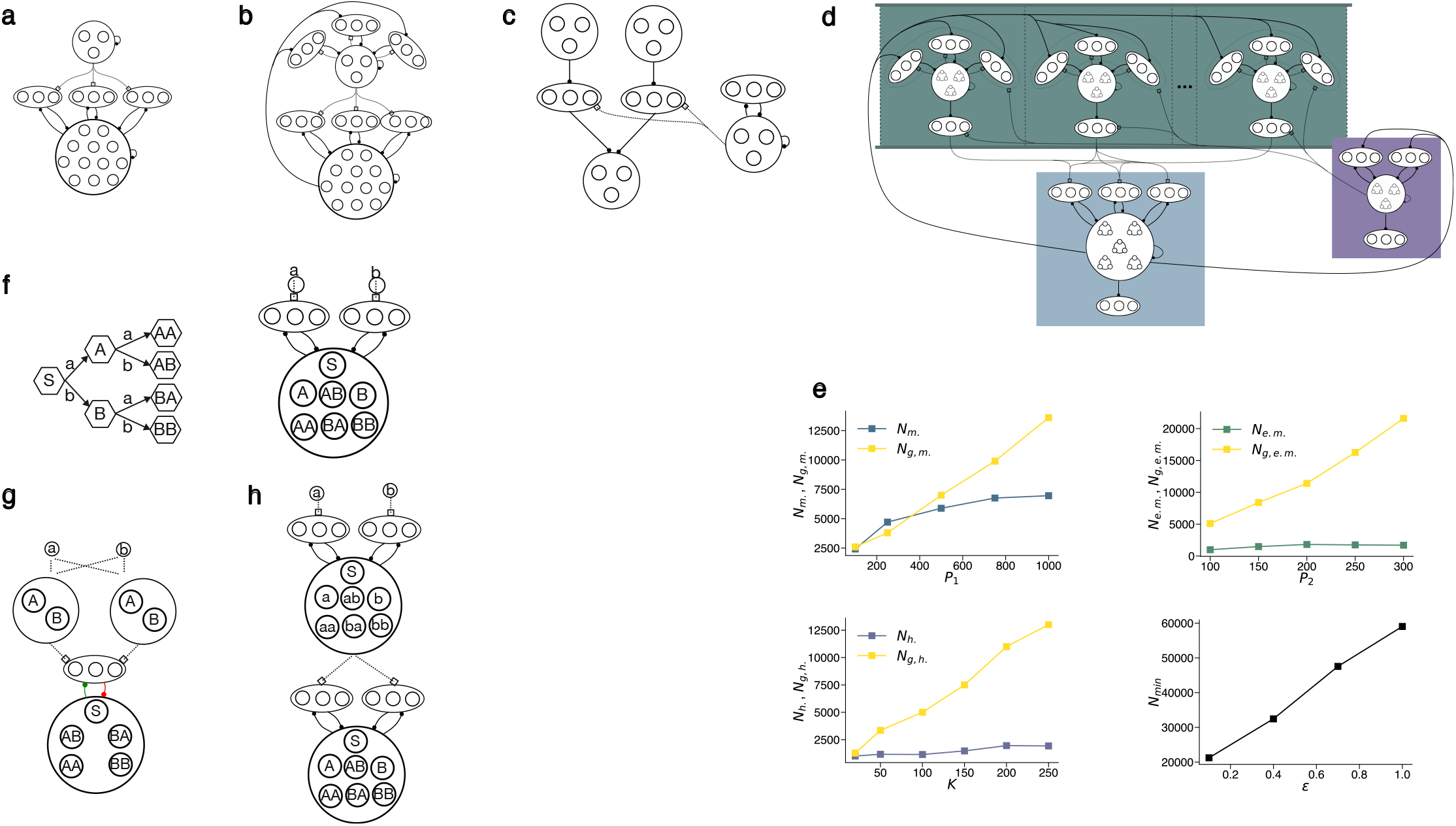
Assembling neural architectures from multiple NLSC. **a**, NLSC augmented with an external memory to produce behaviors with long-distance dependencies. **b**, Two NLSC piled-up, e.g. to produce sequences organized into chunks. **c**, Using NLSC with output gate neurons as memory slots (top), with a conditional routing to the machine NLSC (bottom) by the head NLSC (right), e.g. to produce sequences following an algebraic pattern. **d**, Assembling NLSC to build neural Turing machines. **e**, Number of neurons in machine, external memory, and head circuits when the number of machine states, symbols of the external memory, external memory cells are varied. **f**, FSA for mapping temporal sequences of symbols into meaning states (left) and its NLSC implementation (right). **g**, Neural architecture with an intermediary memory buffer composed of two slots implemented by dictionaries. The machinery to fill slots from sequentially presented inputs is not described, but see circuit Fig. 3b. **h**, Coupling NLSC to map sequences organized into chunks.

